# Goal-directed motor actions drive acetylcholine dynamics in sensory cortex

**DOI:** 10.1101/2021.12.21.473699

**Authors:** Jing Zou, Jan Willem de Gee, Zakir Mridha, Simon Trinh, Andrew Erskine, Miao Jing, Jennifer Yao, Stefanie Walker, Yulong Li, Matthew McGinley, Samuel Andrew Hires

## Abstract

Numerous cognitive functions including attention and learning are influenced by the dynamic patterns of acetylcholine release across the brain. How acetylcholine mediates these functions in cortex remains unclear, as the relationship between cortical acetylcholine and behavioral events has not been precisely measured across task learning. To dissect this relationship, we quantified motor behavior and sub-second acetylcholine dynamics in primary somatosensory and auditory cortex during rewarded sensory detection and discrimination tasks. We found that acetylcholine dynamics were directly attributable to goal-directed actions (whisker motion and licking), rather than delivery of sensory cues or rewards. As task performance improved across training, acetylcholine release associated with the first lick in a trial was strongly and specifically potentiated. These results show that acetylcholine dynamics in sensory cortex are driven by directed motor actions to gather information and act upon it.

## Introduction

Acetylcholine is a major neuromodulator in the brain that influences diverse cognitive functions that span timescales, including arousal ^1, 2, 3, 4^, selective attention ^5, 6, 7, 8^, reinforcement learning ^1, 9^**^;^** ^10, 11, 12^ and neural plasticity ^13, 14, 15, 16^. Many of these functions are mediated through acetylcholine’s influence on cortical circuits ^17, 18^. Cholinergic projections to cortex are complex, arising from multiple basal forebrain (BF) nuclei that contain neuronal subgroups with distinct projection specificity and arbor distributions within and across projection areas ^3, 19, 20^. Individual nuclei also show significant differences in the behavioral events which correlate with their activity patterns ^3, 21^. Thus, the full array of cholinergic dynamics in projection targets cannot be inferred from observations of a single originating area. Direct observation of the spatiotemporal dynamics of acetylcholine in cortex bypasses this organizational complexity and can provide insight to how convergent cholinergic inputs influence cortically regulated cognitive functions.

Numerous lines of evidence demonstrate that the cortical release of acetylcholine regulates arousal and attention ^17, 18^. Increased cortical acetylcholine levels are associated with ^22^ and required for induction of cortical desynchronization during active sensing ^23^. Acetylcholine release causes layer-specific modulation of responses in primary somatosensory cortex (S1) to whisker stimulation ^5, 24^, enhances sensory evoked responses in primary auditory (A1) ^7^ and visual (V1) cortex ^25, 26^, and suppresses spontaneous activity in S1 during whisker movement ^23^. These effects improve stimulus discriminability and provide a mechanism for selective attention.

Attention is crucial for learning and performing tasks. Acetylcholine’s role in these processes is becoming better appreciated through accumulating studies identifying which cholinergic neurons fire at what times during task acquisition and execution. Cholinergic neurons in BF respond ^11^ and cortical acetylcholine transients are evoked to reinforcement-predictive sensory cues ^1^. The extent to which association learning sculpts these responses varies across reports, with such stimuli driving increasing amounts of cholinergic activity in BF (tones and punishment ^16^, odors and reward ^12^) and in nucleus basalis (NB) to basolateral amygdala (BLA) projections (tones and reward ^11^) across training, contrasting with the finding that reward-predictive tones show stable acetylcholine release in BF with learning ^21^. Cholinergic neurons throughout BF also strongly respond to negative reinforcement ^12, 16, 21, 27, 28^, but are inconsistently reported to respond to positive reinforcers like reward. In primates, 70% of all BF neurons were significantly modulated in a choice period, but only 25% in a reward period ^29^. Cell-type specific recordings in rodents found cholinergic neurons within BF do respond to positive reinforcers, scaled by reinforcement surprise ^27^ and encode valence-free reinforcement error ^12^. However, in rat medial prefrontal cortex (mPFC), acetylcholine levels increased for reward-predictive cues, but not for reward delivery ^1^.

A challenge in interpreting studies of cholinergic activity is that both reinforcement-predictive cues and reinforcing events are tightly correlated with orofacial and body movements ^30, 31^. These movements are also associated with transient increases in cortical acetylcholine levels, which suggests acetylcholine may link action and expectation ^4, 32, 33^. Of particular interest to reward related mechanisms, licking is reported to drive cholinergic activity, though reports vary from strong acetylcholine release at lick bout onset ^28, 34^ and offset ^21^, to weak release to licks ^12^ to none at all ^27^. This variability may arise from experimental differences in task conditions and sensory modalities or may reflect a compartmentalization of acetylcholine functions based on nucleus, cell-type, and projection targets. Regardless, since whisking and licking patterns are strongly influenced by reward expectation and delivery ^30,35^, dissociating sensory cues and reward from these motor actions is crucial for interpreting acetylcholine’s role in task learning and performance.

Here we sought to constrain how acetylcholine influences sensory cortex by measuring spatiotemporal dynamics of acetylcholine in S1 and A1, areas that undergo remarkable representational plasticity during learning ^36^. We directly imaged sub-second changes in acetylcholine concentration using a GPCR Activation Based sensor (GRAB-ACh3.0 ^32^) broadly expressed on the surface of supragranular cortical neurons. High-speed videography during active whisker-guided object location discrimination (S1) and sound detection (A1) allows dissociation of cue and stimulus from tightly coupled motor responses, while our response design allows dissociation of reward delivery from the motor action of licking. Recording across weeks of training, we identify behavioral correlates of acetylcholine release in sensory cortices during task performance and how tactile discrimination learning specifically reorganizes those dynamics in S1 across the transition from novice to master.

## Results

### Mice can learn to discriminate object location using a single whisker

To measure the dynamics of acetylcholine in primary somatosensory cortex during tactile discrimination learning, we employed a go/no-go single-whisker object localization task (**Figure 1A** ^37^). We trained water-restricted head-fixed mice (n=8 mice) to search with a whisker for the position of a thin steel pole presented in one of two locations along an anteroposterior axis during a one second sampling period (**Figure 1B**), and to lick for a water reward during a subsequent one second answer period if the pole was in the posterior location. Mice were cued to the presentation and removal of the pole by the sound of a pneumatic valve. Licking during the sampling period had no consequences, while licking in the answer period determined trial outcome and extended the duration of the pole presentation.

**Figure 1.**
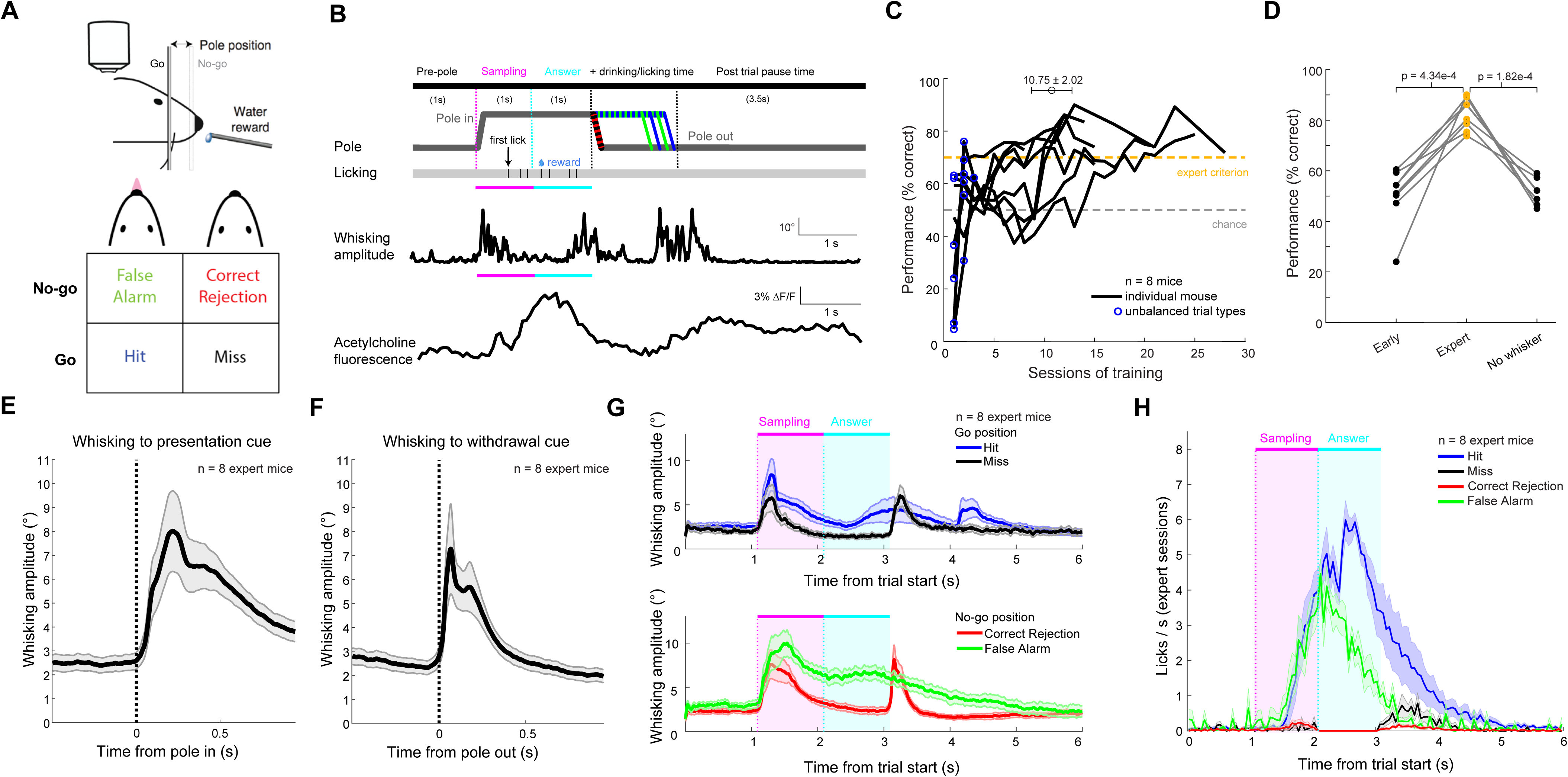
Learning of whisker-guided object location discrimination and associated motor actions. **A.**Two-location discrimination task design with trial outcomes. **B.** Single trial structure with example licking, whisking, GRAB-ACh signal traces. **C.** Learning curves of 8 mice, mean ± SEM sessions to expert. Blue circles, unbalanced go and no-go trials. Gold dash, expert threshold. Gray dash, chance level. **D.** Mean performance for three early and expert sessions, per mouse. 3 mice with 1 no-whisker session, 4 mice with 3 no-whisker sessions averaged. **E.** Average whisking amplitude aligned to pole in cue. Mean and SEM. 3 expert sessions / mouse. **F.** Same aligned to pole out cue. **G.** Grand mean trial averaged whisking amplitude by trial outcome. Mean and SEM. 3 expert sessions / mouse. **H.** Same for licking rates.

Mice were trimmed to their C2 whisker at least one week prior to onset of imaging experiments. All mice reached task mastery (>70% performance for 3 consecutive sessions) within 1-3 weeks of training (mean 10.75 ± 2.02 sessions; mean 3373.75 ± 754.09 (standard error) trials **Figure 1C**). Mean Hit rates rapidly increased within three sessions of training, while mean False Alarm rates slowly declined across weeks (**SFigure 1A**). Trimming of the C2 whisker after task mastery dropped performance to chance (**Figure 1D**), demonstrating the whisker-dependence of the task. Expert mice initiated whisker exploration upon the pole presentation and withdrawal cue sounds (**Figure 1E, F**, **SFigure 1B**). Since whisking amplitude was physically restricted when the pole was in the go-associated proximal position, we examined the whisking patterns of go trials vs. no-go trials separately. For both positions, trials with licks during the sampling and answer periods (i.e. Hit and False Alarm) had more sustained whisking than trials without licks (i.e. Miss and Correct Rejection; **Figure 1G**). This difference is consistent with licking-coupled whisker motion ^38^. Lick rates on Hit and False Alarm trials were indistinguishable during the sampling period and diverged during the answer period once water was dispensed due to reward collection on the Hit trials (**Figure 1H**). Licking rates on Correct Rejection and Miss trials were zero by construction during the answer period, with occasional licks outside this period.

### Acetylcholine release patterns differentiate trial types

Acetylcholine dynamics were measured using two-photon imaging of supragranular layers of S1. The C2 whisker barrel was targeted for viral injection of AAV9-GRAB-ACh3.0 via intrinsic signal imaging (**SFigure 2A**). Behavioral training commenced three weeks post injection, when strong fluorescence signals were present in all mice (**Figure 2A**). To minimize the impact of a rapid partial dimming of fluorescence following illumination onset (**SFigure 2B**), two-photon imaging and illumination (15.44 fps, 940nm, 30-60mW out of objective) was continuous from the session start, and the first 100 seconds of the session were excluded from analysis. Phasic increases in fluorescence across the entire field of view were observed shortly after most, but not all onsets of pole presentation (**Figure 2B**). These fluorescence dynamics diverged across trial outcomes (**Figure 2C**). Hit and False Alarm trials showed a strong, similar increase in signal during the sampling and answer periods (example session, **SFigure 2C**; grand mean of expert sessions **Figure 2D**). This was followed by a secondary response that, on average, persisted into the inter-trial period. In contrast, Miss and Correct Rejection trials on average showed two short latency, short duration increases in fluorescence following pole presentation and withdrawal. We did not find evidence for differential regulation of acetylcholine dynamics at the spatial scale of cortical columns, as trial type averages within the primary whisker column versus in the surrounding columns were identical for all trial types (**Figure 2E**).

**Figure 2.**
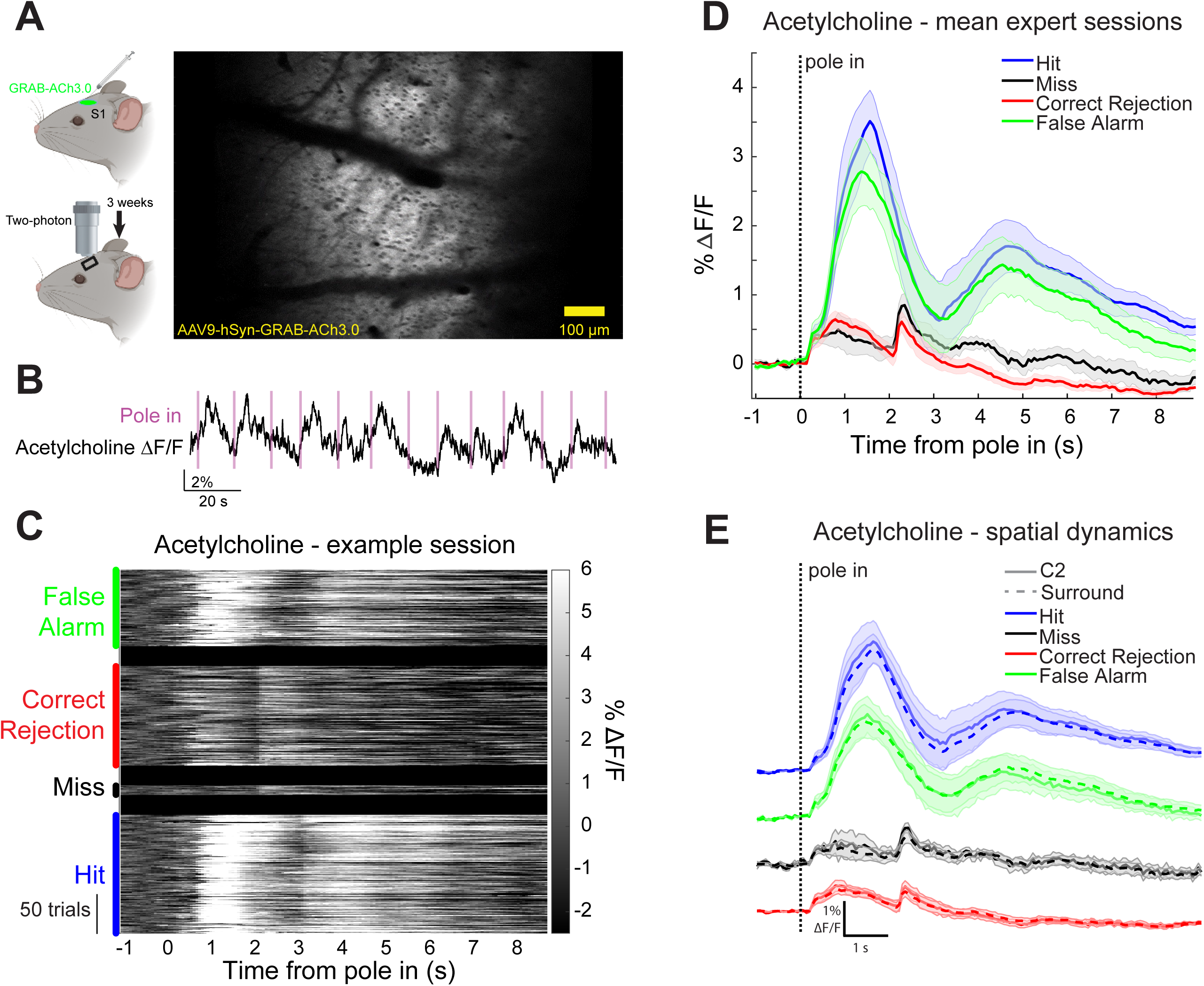
Acetylcholine release in S1 varies with trial outcome. **A.** GRAB-ACh3.0 AAV expression in S1 barrel cortex, 3 weeks post injection. Cartoon generated by BioRender.com. **B.** Phasic increase of acetylcholine release after pole in cue in most trials. **C.** Acetylcholine induced fluorescence changes sorted by trial types, one expert example session. **D.** Grand average, 8 mice, 3 expert sessions each. **E.** Comparison of grand average acetylcholine dynamics within the C2 column (solid) and in the surround (dashed). 8 mice, 3 expert sessions each. Traces are vertically offset for display.

### Whisking and licking but not reward induce acetylcholine release in sensory cortex

To investigate potential triggers of acetylcholine release in cortex, we compared the acetylcholine dynamics across trial types to several classes of behavioral events, including pole presentation and withdrawal cues, whisker exploration, whisker touch, licking, and reward. All trial types shared a common time for pole presentation, and all trial types showed a small, short latency (2-3 frames, 130-195ms) increase in acetylcholine aligned to that cue (**Figure 3A**). Lick trials showed much bigger and longer transients immediately following this initial hump. Trials without licks in the answer period had a common pole withdrawal time, while trials with answer licks had a variable withdrawal time. Alignment to pole withdrawal again showed a sharp upward transient of acetylcholine across all trial types, though this was overlaid on longer duration acetylcholine dynamics (**Figure 3B**). In all cases, the initial rise in fluorescence began within 1-2 frames after a sharp increase in whisking amplitude (**Figure 3A,B**).

**Figure 3.**
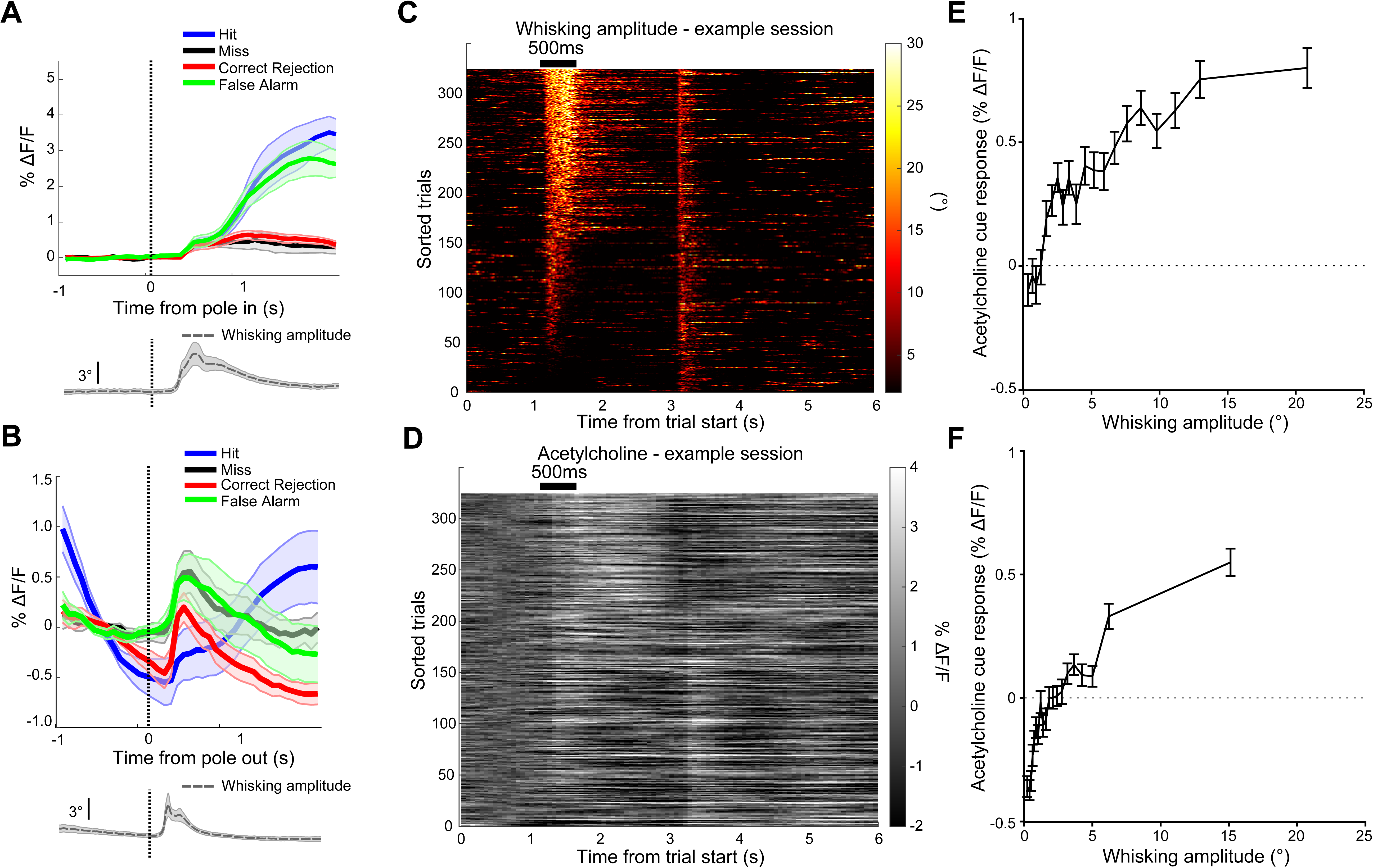
Whisking drives acetylcholine release in S1. **A.** Top, grand mean acetylcholine fluorescence change aligned to pole in cue. Bottom, average whisking amplitude. 8 mice, 3 expert sessions each. **B.** Same to pole out cue. **C.** Whisking amplitude sorted by mean of 500ms post pole in cue. No lick trials pooled from 3 expert sessions, 1 mouse. **D.** Same, for acetylcholine fluorescence change sorted by whisking amplitude. **E.** Grand mean of acetylcholine fluorescence change vs. whisking amplitude of 500ms after pole in cue. No lick trials from 3 early and 3 expert sessions, 8 mice. **F.** Grand mean of acetylcholine fluorescence change vs. whisking amplitude during 1s before pole in cue for expert sessions. Trials with no licks within 1.5s after trial start, 3 expert sessions, 8 mice.

Was acetylcholine release driven by the pole presentation cue *per se*, or was it driven by a motor response to the cue (**Figure 1E,F**)? To test this, we examined the covariation of cue-evoked whisking with cue-associated acetylcholine dynamics. We restricted our analysis to no-lick trials to avoid potential confounds of licking-driven acetylcholine responses. We sorted acetylcholine responses in no-lick trials by the average amplitude of the whisker motion within 500 milliseconds after the pole presentation cue (**Figure 3C, D**). A fraction of trials (23.9% mean ± 21.8% per mouse) did not evoke whisker motion (<2° mean amplitude) upon pole presentation, with most producing a range of whisking vigor (**SFigure 3A**). In trials without cue-evoked whisking, there was no acetylcholine release following the cue (0.01% mean ΔF/F for whisking amplitude <2°), while trials with cue-evoked whisking showed a positive relationship between whisking amplitude and acetylcholine response (0.58% mean ΔF/F for whisking amplitude >5°; 0.08% mean increase in ΔF/F per degree of amplitude (**Figure 3E**, **SFigure 3B**). These findings were recapitulated in pole withdrawal-cued whisking and acetylcholine responses (**SFigure 3C-E**). This implies that whisking, rather than whisker-pole contact drove the responses, since whisking after pole withdrawal rarely causes pole touches. Finally, whisking bouts prior to the presentation cue showed a similar positive relationship between whisking amplitude and acetylcholine response (0.0642% mean increase in ΔF/F per degree of amplitude), with a more negative fluorescence offset in the non-whisking condition attributable to an earlier baseline period (**Figure 3F**). These data demonstrate that cue-associated acetylcholine release in S1 is directly proportional to the magnitude of the motor response (i.e. exploratory whisking) evoked by that cue, and this motor relationship is maintained in the absence of cue. We conclude that motor actions, rather than the sensory cue, drives cue-associated acetylcholine transients in S1.

The striking difference in acetylcholine dynamics across trial types (**Figure 2C-E**), in particular Hit and False Alarm versus Miss and Correct Rejection, suggested that licking could be a powerful driver of acetylcholine release in S1. We aligned trials across expert sessions to the time of first lick, regardless of trial type or if the lick occurred in the answer period, and sorted by the number of licks in the trial (**Figure 4A**). Trials with no licks were aligned to the time of the median first lick. Licks were tightly clustered following the first lick (**Figure 4B**) and preceded rhythmically at a regular modal inter-lick interval of 155ms in expert mice (**Figure 4C**). On trials with licks, there was a sharp increase in acetylcholine signal beginning shortly before lickport detection of the first lick, which lasted 1-2 seconds (**Figure 4D**). This lick-associated response dwarfed the cue-associated transients on the no-lick trials (**Figures 2E)**, and partially scaled with the number of licks **(Figure 4D)**. Across all expert mice, the first lick was associated with a profound increase in mean acetylcholine over the following second, jumping from 0.35% ± 0.26% on no-lick trials to 1.60% ± 0.42% with a single lick (**Figure 4E**), an increase of 1.25% ΔF/F over baseline. Subsequent licks yielded smaller increases in mean acetylcholine over this period at a rate of 0.28% ΔF/F per additional lick. Similarly, the duration of the transient was 1.46 ± 0.65s for a single lick. Subsequent licks extended this transient 116ms per lick (**Figure 4F**), somewhat less than the inter-lick interval (**Figure 4C**). This demonstrates that acetylcholine release in S1 is primarily coupled to the onset of goal-directed licking, with the vigor of licking modulating the magnitude and duration of the release.

**Figure 4.**
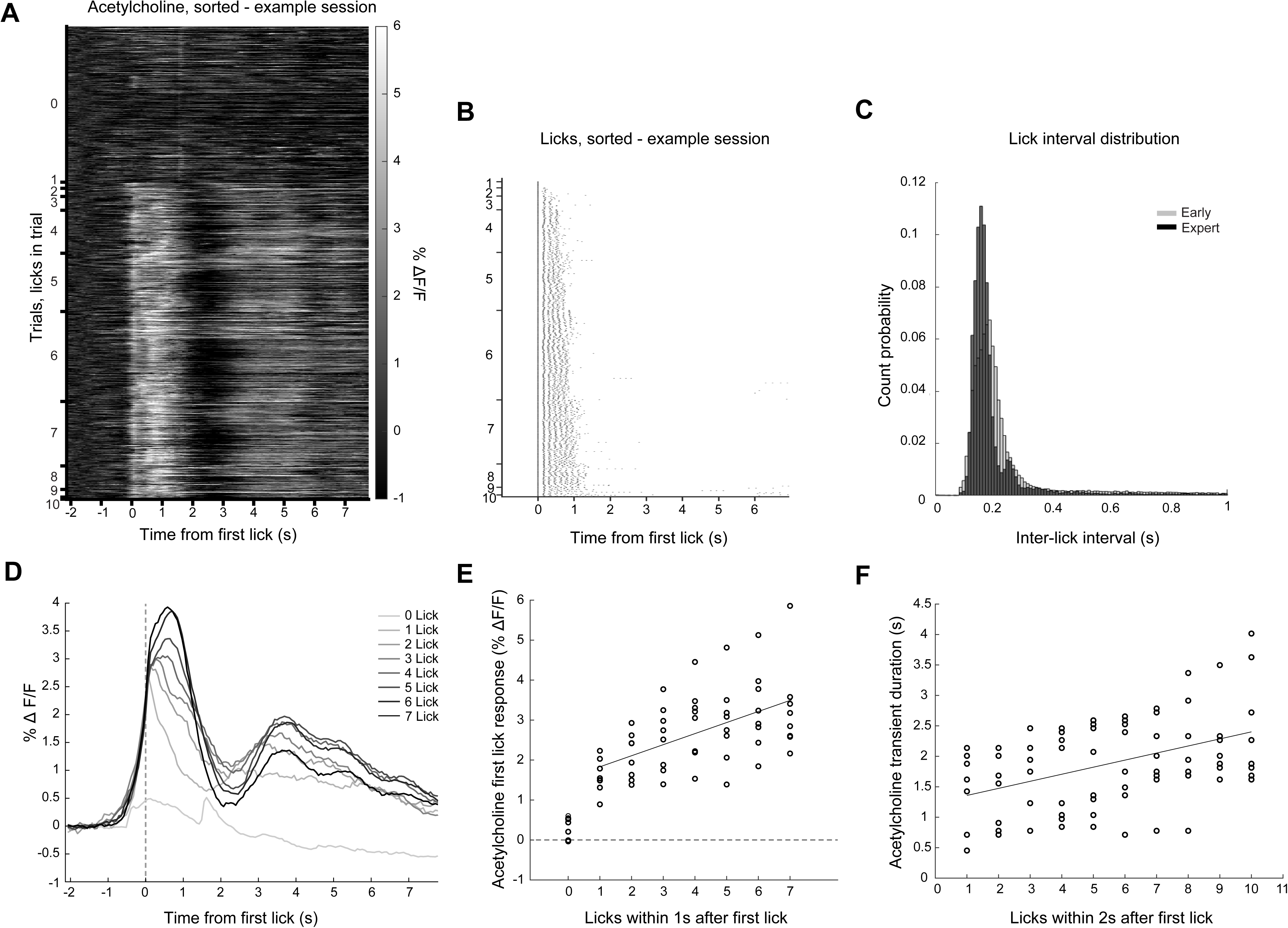
Licking strongly drives acetylcholine release in S1. **A.** Acetylcholine fluorescence change across trials, sorted by number of licks in trial, aligned to first lick. One example expert session. **B.** Lick pattern across trials, sorted by number of licks in trial, aligned to first lick. Same example session as A. **C.** Pooled inter-lick interval from 3 early (gray) and 3 expert (black) sessions each, 8 mice. **D.** Acetylcholine fluorescence change aligned to first lick for trials with different numbers of licks. Same session as A. **E.** Mean acetylcholine fluorescence change in 1 second following first lick binned by number of licks in that period. Circles, mean 3 expert sessions / mouse. Linear fit from 1-7 licks. **F.** Duration, onset to trough, of the first acetylcholine transient, binned by number of licks within 2 seconds of first lick. Circles, mean 3 expert sessions / mouse. Linear fit from 1-10 licks.

During the period of goal-directed licking (0-1s post-first lick), there was a small, but significant increase on Hits over False Alarm trials (**Figure 5A**, **SFigure 4A**). Could this difference be explained by reward delivery? Rewards are only distributed upon a correct lick in the answer period. Hit trials had higher acetylcholine levels than False Alarms during the answer period, but also had more licks (**Figure 5B,C**). To control for increased acetylcholine release caused by additional licking (**Figure 4E**), we pairwise compared mean acetylcholine levels of Hit and False Alarm trials during the answer period for matched numbers of licks in the period. Hit and False Alarm acetylcholine levels were essentially identical after accounting for the difference in lick count (**Figure 5D**). There was no difference in the distribution of first lick times of Hits and False Alarms (**SFigure 4B**). We conclude that reward delivery does not induce acetylcholine release in supragranular S1, consistent with prior electrochemical measurements in mPFC ^1^. Together, these data establish that the main driver of acetylcholine release in supragranular S1 during tactile-guided choice behavior is execution of the goal-directed action, secondary drivers are exploratory whisker motion and additional licking, while task initiation cues and reward delivery have no direct effect.

**Figure 5.**
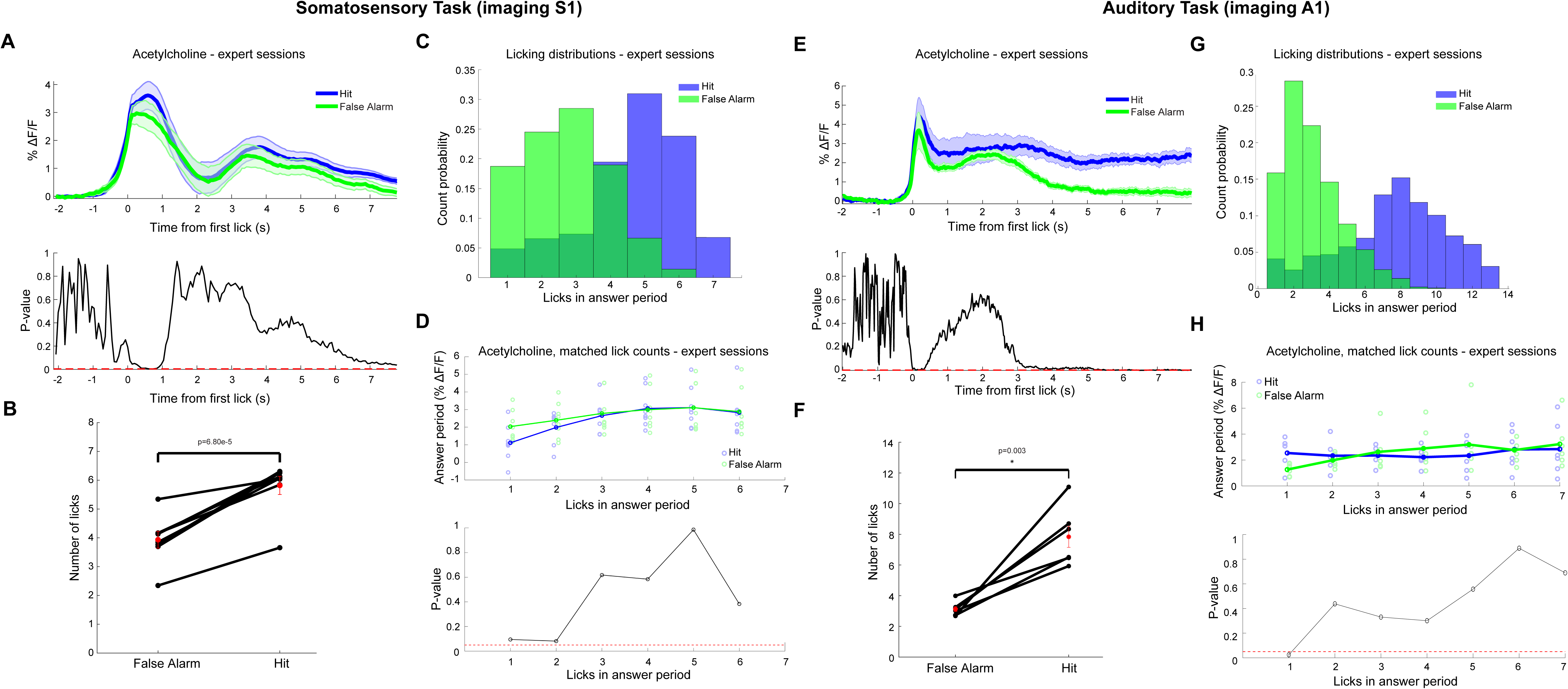
Reward delivery does not drive acetylcholine release in sensory cortex. **A.**Top: Acetylcholine fluorescence change aligned to first lick from Hit (blue) and False Alarm (green) trials. Mean and SEM. 3 expert sessions, 8 mice for all panels. Bottom: Significance test between Hit and False Alarm acetylcholine over time (p-value, paired t-test, corrected for multiple comparisons). **B.** Mean number of licks in the answer period on Hit and False Alarm trials. **C.** Distribution of lick counts in the answer period histogram for Hit (blue) and False Alarm (green) trials. **D.**Top: Mean acetylcholine fluorescence change in answer period, binned by licks in answer period for Hit (blue) and False Alarm (green). Bottom: Significance of difference between Hit and False Alarm trials, binned by licks (p-value, paired t-test, corrected for multiple comparisons). **E-H.** Same as A-D, but in auditory cortex .

To investigate whether the above relationship between licking, reward delivery and acetylcholine dynamics generalizes across sensory cortical areas, we performed an auditory detection task while imaging GRAB-ACh3.0 responses in primary auditory cortex. As in S1, licking drove strong phasic increases of Ach signal, with Hit trials showing higher peak and secondary response magnitude than False Alarms (4.44% vs 3.69%; **Figure 5E**, **SFigure 4C**). However, Hit trials had more licks (**Figure 5F, G**) than False Alarms. Pairwise comparison of Ach signals in lick count matched trials showed no significant difference between Hits and False Alarms (**Figure 5H**). This reinforces our conclusion that reward-seeking licking drives acetylcholine release across sensory cortex, while reward delivery does not.

### Mastering the task enhances first lick induced acetylcholine release

Acetylcholine regulates attention ^39^, which may change with task performance, familiarity, or associative learning. It follows that training might increase acetylcholine release concurrently with improved performance. Furthermore, learning to associate sensory stimuli with potential reward increases cholinergic neuron activity in the basal forebrain during stimulus presentation ^12^. On the other hand, training might reduce acetylcholine release, as a familiar task may require less attentional resources to solve or reduced neural plasticity once established. To address these possibilities, we compared the acetylcholine dynamics for the first three sessions of the full somatosensory task training and the final three expert sessions in each mouse. We found that the initial lick-associated acetylcholine release was nearly twice as large in expert sessions compared to early training sessions (ΔF/F 3.07 ± 1.01% std expert vs. 1.67 ± 0.65% std early; **Figure 6A**). The magnitude of the increase was directly proportional to session performance with mean ΔF/F increasing 0.53% per 10% increase in correct rate (**Figure 6B**). This increase occurred even though expert mice licked fewer times (**Figure 6C,D**) and at the same pace (**Figure 6E**) as compared to early sessions. With training, expert mice consolidated their licking into the sampling and reward windows, which follow the cue presentation (**SFigure 5A**). Restricting our analysis to only trials where the first lick came after the cue confirmed that acetylcholine responses associated with the first lick are significantly potentiated after training (**SFigure 5B**).

**Figure 6.**
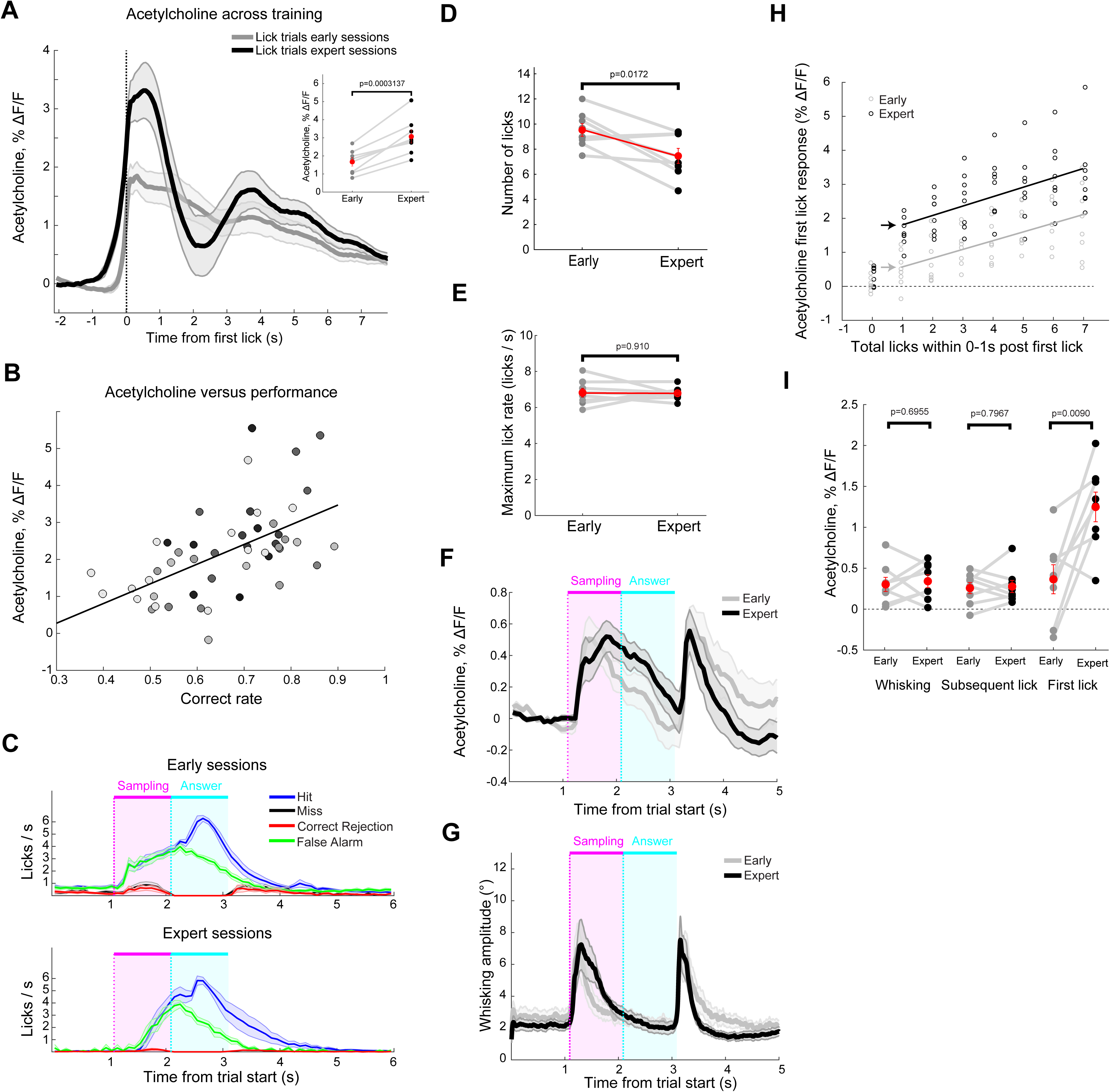
Learning selectively potentiates acetylcholine release to first licks in S1. **A.** Grand mean acetylcholine fluorescence change aligned to first lick in early (gray) and expert (black) sessions, 3 sessions per condition, 8 mice for all panels except Paired comparison of average acetylcholine fluorescence change within one second after first lick between early (gray) and expert (black) sessions for each individual mouse is shown in inset figure. **B.** Bands SEM. B. Relationship between correct answer rate and acetylcholine fluorescence change within 1 second following first lick. Shade indicates mouse identity. All 51 single-whisker imaged sessions, Fit equation r = 0.05328*x-0.0123. **C.** Mean lick rates for trial types in early (top) and expert (bottom) sessions. **D.** Mean lick numbers per trial. **E.** Peak lick rate. **F.** Grand mean ± SEM acetylcholine fluorescence change in no lick trials. **G.** Same for mean ± SEM average whisking amplitude **H.** Mean acetylcholine fluorescence change in 1 second following first lick binned by number of licks in that period. Linear fits from 1-7 licks. **I.** Acetylcholine fluorescence change from whisking, subsequent lick, and first lick for early (gray) and expert (black) sessions. Red, grand mean ± SEM.

This increase of acetylcholine signal after training was not due to an increase of GRAB-ACh sensor expression or sensitivity seen by the following internal control. On no-lick trials, whisking-associated acetylcholine release slightly increased following the pole-presentation cue, and slightly decreased following the pole-withdrawal cue (**Figure 6F**). This closely matched the change in whisking to those cues following training (**Figure 6G**). Thus, the relationship between acetylcholine fluorescence change and whisker motion was stable across training. Finally, we examined whether training induced potentiation of all licks, or only the first lick (i.e. the choice-signaling lick in expert sessions), by repeating the analysis of **Figure 4** on early sessions of the same cohort of mice. In contrast to experts (**Figure 4E**), the first lick in early sessions drove a modest increase in acetylcholine from 0.14% ± 0.3% on no-lick trials to 0.50% ± 0.52% with a single lick (**Figure 6H**), an increase of 0.36% ΔF/F over baseline. Subsequent licks caused a 0.26% increase in ΔF/F per additional lick versus 0.28% ΔF/F per lick in expert mice (**Figure 6H**). Thus, training induced a nearly 3.5x increase in acetylcholine response over baseline specifically to the first lick, while leaving responses to whisking and subsequent licks unchanged (**Figure 6I**). We conclude that training induces a dramatic and selective potentiation of acetylcholine release in supragranular S1 to a goal-seeking motor action and this potentiation is correlated to improved task performance.

### First lick is the major predictor of acetylcholine release

Our above analyses identified heterogenous influences of behavioral events with overlapping time distributions on acetylcholine dynamics in sensory cortex. To further tease apart these interactions we trained a general linear model (GLM) to predict cholinergic dynamics in S1 from the recorded events. The predictors used included sensory stimuli (pole in and out auditory cues), motor actions (first lick, other licks, and whisking Hilbert components of amplitude, midpoint and phase), reward delivery, and fluorescence history (to account for indicator decay rate). The GLM output closely followed the observed low-frequency fluorescence dynamics on hold-out trials (**Figure 7A, B**). Following training, the weight of the first lick time significantly increased, and was the single greatest predictor of positive fluorescence responses in the first second following lick (**Figure 7C**). Reward delivery predicted a short-term drop in fluorescence in both early and expert mice. Extending beyond the first second, both first lick and reward kernels have a biphasic positive and negative structure, suggesting additional complexities in the interaction of event-triggered release and tonic cholinergic tone.

**Figure 7.**
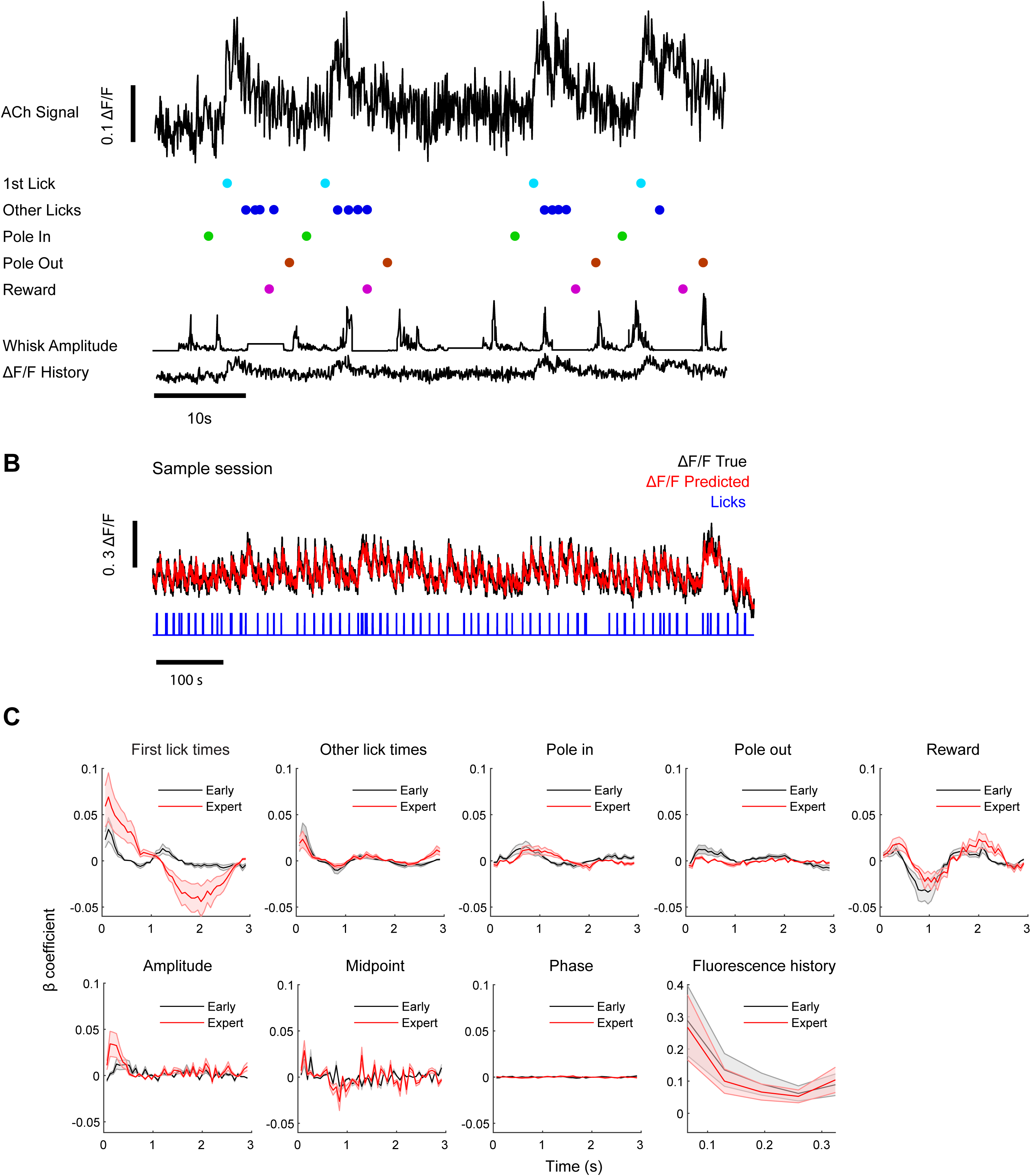
General linear model of acetylcholine fluorescence dynamics from sensory, motor, and reward predictors. **A.** Sample acetylcholine fluorescence trace. Object model fit to predict fluorescence from below predictors. **B.** Sample true fluorescence (black), corresponding predicted fluorescence trace using the predictors in A (red), and licks (blue). **C.** Different predictors’ β coefficient for early (black) and expert (red) sessions. 3 sessions for each condition, 7 mice.

## Discussion

Recording and manipulation of cholinergic neurons originating in BF nuclei ^26, 27, 34, 40, 41^ has established the importance of cholinergic signaling on multiple brain functions. However, the complexity of BF organization ^3, 19, 20^ has posed a challenge in linking specific cognitive functions to acetylcholine dynamics in specific projection targets. By recording acetylcholine dynamics directly in S1 (**Figure 2**) during whisker-guided object localization (**Figure 1**), we discovered surprising differences in the triggers and dynamics between previously observed BF nuclei and this cortical target essential for tactile discrimination ^42^. Nearly all acetylcholine release in supragranular S1 was attributable to goal-directed motor actions (**Figures 3, 4**) rather than sensory input (**Figure 3**) or reward delivery (**Figure 5**). This generalized for licking and reward in a second sensory area (A1, **Figure 5**). Moreover, as task performance improved across training, acetylcholine release to the first lick in a trial became dramatically and specifically potentiated (**Figures 6**, **7**).

Together, these data support a model that acetylcholine release in sensory cortex during reward-seeking behavior is driven by directed motor actions to gather information (e.g. whisking) and act upon it (e.g. licking). This contrasts from reports that acetylcholine release is driven by delivery of reinforcement-predictive cues ^11, 12, 16^ and reward itself ^11, 12^. In contrast to passive cue-reward association, our task requires active exploration to gather reward-predictive information, revealing a motor requirement for acetylcholine release. Intriguingly, cortical VIP neurons, which are strongly excited by acetylcholine ^43^, are activated by motor actions that are associated with the expectation of reward (e.g. licking), but not reward delivery ^44^. In a similar context, our observed potentiation of acetylcholine release to the first lick is consistent with a learned link between reward-expectation and a specific motor action. Our results also suggest that cue-induced changes in motor behavior (e.g. increased whisking, sniffing or anticipatory licking) and other orofacial movements ^33^ may also provide a meaningful contribution to acetylcholine release in passive association tasks.

The lack of reward-evoked acetylcholine release in S1 may be due to several factors. One possibility is that reward activity in BF is transmitted only to particular targets such as BLA ^11^, but not S1, due to projection specificity in subpopulations of BF cholinergic neurons ^3^. Second, while reward-delivery signals have been observed in cholinergic neurons of HDB and NB ^27^, fiber photometry in those areas show reward-associated transients are small relative to lick-evoked transients and tightly correlated with an increase of lick rate following water delivery ^21^. Perhaps reward-delivery only indirectly drives acetylcholine release via changes in licking patterns. However, this result may depend on task and reward structure. Tasks with variable reward probability have shown decreased cholinergic activity to delivery of highly likely rewards ^12, 27^. In our task, licking on go trials guaranteed reward, which may have shifted reward delivery-associated activity earlier to the time of the first lick, when reward is expected.

Intriguingly, acetylcholine release associated with the first lick began ramping several hundred milliseconds prior to lick detection (**Figure 4C**). This may be due to a combination of following factors: whisking precedes licking and drives modest and additive acetylcholine release (**Figure 3**), decision related activity precedes motor action by some amount of time ^45^, and the tongue requires about 150-200ms from protrusion initiation to lickport contact ^35^. These are partially counterbalanced by the indicator rise time, which is dependent on the underlying acetylcholine concentration profile and likely on the order of tens of milliseconds for a few percent ΔF/F change ^32^. Thus, some of the first lick-associated acetylcholine dynamics may be caused by an internal choice deliberation associated with initiation of motor action. This possibility is reinforced by the observation that onset of acetylcholine release is more closely aligned to first lick on early than on expert sessions (**Figure 6A**), suggestive of expert-specific choice-associated dynamics overlaid on motor-triggered dynamics common to both session classes.

The patterns of acetylcholine release in response to whisking and licking suggest potential functions for acetylcholine on sensory cortical circuitry ^46, 47^. VIP cells in S1 are activated by whisking ^48^ via acetylcholine release ^49^ leading to disinhibition of excitatory neuron dendrites where top-down contextual information arrives in S1 ^50, 51, 52^. Thus, whisking-induced acetylcholine release could enhance integration of contextual information with sensory input in S1 neurons, providing, for example, a potential mechanism for combining motor and touch signals to generate location specific codes ^53^ and percepts ^52^. VIP activation also gates neural plasticity in cortex ^55^. Higher task performance is associated with increased acetylcholine in V1 ^26^. Our similar results in S1 (**Figure 6B**) identified that this increase is specific to the first lick and persists for several seconds (**Figure 4**). Thus, this increase is well-poised to provide windows of enhanced neural plasticity via VIP cell activation while the consequences of decisions are evaluated.

This work is only a step towards understanding the function of acetylcholine dynamics in cortex and sensory processing. The specificity of acetylcholine action on particular cortical cell types raises the question of the extent to which this reflects differing patterns of receptor expression ^56^ versus preferential targeting by cholinergic axons. While we did not see a difference in acetylcholine dynamics between center and surrounding cortical columns, there could be substantial heterogeneity in release at cellular and subcellular scales. The GRAB sensor expression across the plasma membrane of all neuron types coupled with the intrinsic sensor kinetics made our observations well-suited for quantification of volume transmission, but could obscure sites of fast synaptic transmission. We also did not determine the extent to which the S1 acetylcholine dynamics reflect activity from nucleus basalis (NB) versus horizontal diagonal band (HDB) or their subdivisions which project to S1 ^3^. Cholinergic terminals in mouse neocortex show laminar preferences, entering in either layer 1 or layer 6 depending on the location of originating soma in the BF ^57^. The deeper pathway could potentially convey different classes of information and serve a distinct function from the superficial acetylcholine dynamics observed here. An intriguing possibility is that the selective potentiation of choice-signaling action may be dissociable by afferent source. Finally, while we quantified the triggers and timescales of acetylcholine dynamics on cortical targets, substantial additional work is required to determine the functional consequences of those time limits. Manipulation of local acetylcholine dynamics and cellular targets at specific moments during task performance and acquisition could clarify acetylcholine’s potential roles in regulation of sensory integration and cortical plasticity.

## Acknowledgements

This work was funded by grants U01NS120824 and R01NS102808 from NINDS.

## Author contributions

J.Z., M.J.M. and S.A.H. designed the study. J.Z. and J.W.G. performed the experiments. J.Z., J.W.G., Z.M, S.T., A.E., J.Y. and S.W. analyzed the data and made the figures. M.J. and Y.L.L. provided reagents. All authors reviewed the manuscript. J.Z. and S.A.H. wrote the manuscript.

## Materials and Methods

### Key Resources Table

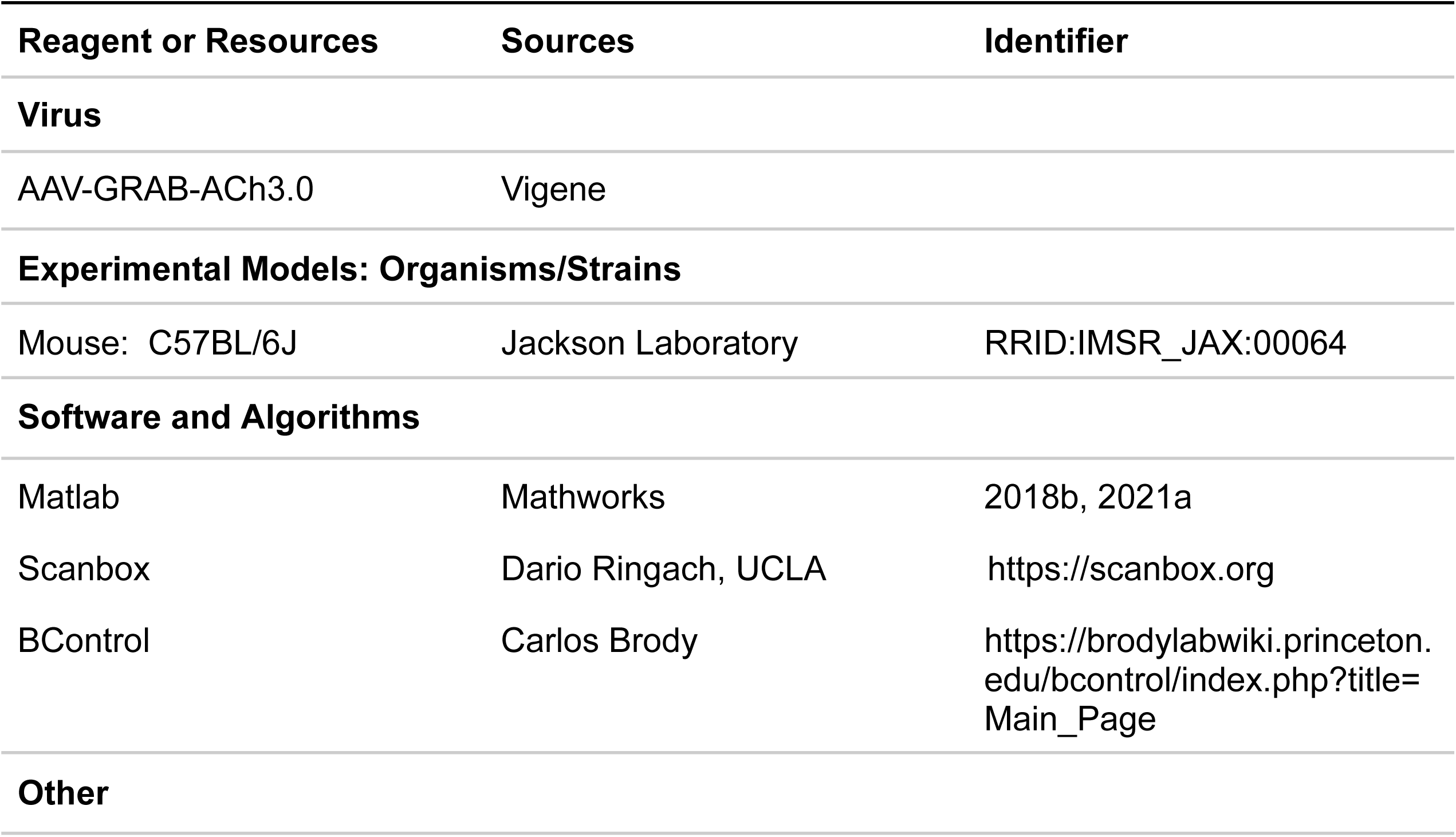
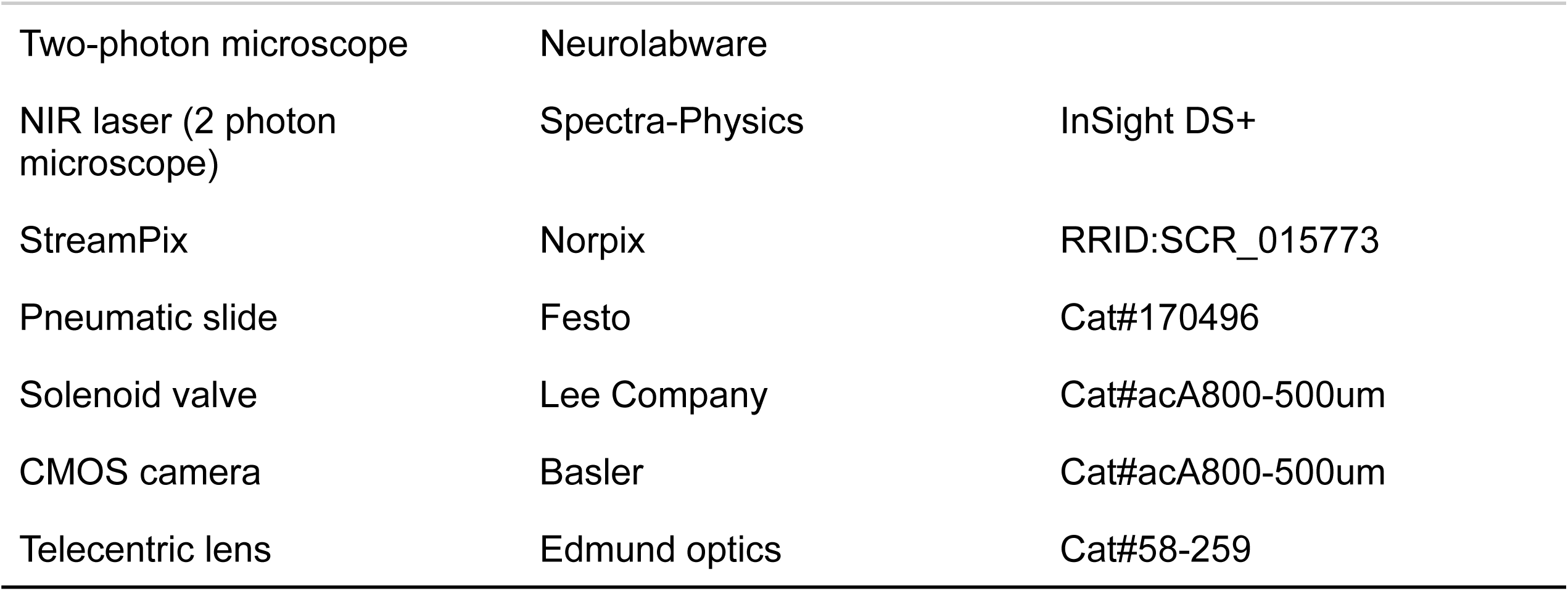

| Reagent or Resources | Sources | Identifier |
| --- | --- | --- |
| **Virus** |  |  |
| AAV-GRAB-ACh3.0 | Vigene |  |
| **Experimental Models: Organisms/Strains** |  |  |
| Mouse: C57BL/6J | Jackson Laboratory | RRID:IMSR_JAX:00064 |
| **Software and Algorithms** |  |  |
| Matlab | Mathworks | 2018b, 2021a |
| Scanbox | Dario Ringach, UCLA | https://scanbox.org |
| BControl | Carlos Brody | https://brodylabwiki.princeton.edu/bcontrol/index.php?title=Main_Page |
| **Other** |  |  |
| Two-photon microscope | Neurolabware |  |
| NIR laser (2 photon microscope) | Spectra-Physics | InSight DS+ |
| StreamPix | Norpix | RRID:SCR_015773 |
| Pneumatic slide | Festo | Cat#170496 |
| Solenoid valve | Lee Company | Cat#acA800-500um |
| CMOS camera | Basler | Cat#acA800-500um |
| Telecentric lens | Edmund optics | Cat#58-259 |

### Lead Contact and Materials Availability

Further information and requests for resources and reagents should be directed to and will be fulfilled by the Lead Contact, Samuel Andrew Hires. This study did not generate new reagents or mouselines

### Experimental Model and Subject Details Animals

For the somatosensory task, we used 2.5-4 months old male (n=2) and female (n=6) C57BL/6J mice (#000664, The Jackson Laboratory). Mice were maintained on a 12:12 reversed light-dark cycle, and put on water restriction 2 weeks before training started. During the water restriction period, health status was assessed everyday following a previously reported guideline^37^. All procedures were performed in accordance with the University of Southern California Institutional Animal Care and Use Committee Protocol 20732 and 20788.

For the auditory task, we used 5 male C57BL/6J mice (The Jackson Laboratory). All surgical and animal handling procedures were conducted in compliance with the ethical guidelines of the National Institutes of Health and received approval from the Institutional Animal Care and Use Committee (IACUC) Protocol AN7369 at Baylor College of Medicine.

## Method Details

### Headbar surgeries

Prior to each surgical procedure, the surgical station and instruments underwent sterilization. Anesthetic gas, specifically isoflurane (2–3% in oxygen), was administered for the entire duration of all surgeries. The mouse’s body temperature was maintained between 36.5°C and 37.5°C using a homoeothermic blanket system.

#### Somatosensory task

Before each surgery, a rimadyl tablet was given 0.5 mg/tablet 24 hours before surgery. buprenorphine-SR and marcaine were injected subcutaneously at 0.5 mg/kg and 2% right before the surgery. A customized stainless steel headbar was attached to the skull using layers of Krazy glue (Elmer’s Products, Inc) and dental cement (Lang Dental Mfg. Co., Inc).

#### Auditory task

Subcutaneous injection of Buprenorphine-SR at a dose of 1.0 mg/kg was administered 30 minutes before anesthetic induction. Once anesthetized, the mouse was positioned in the stereotaxic frame. The surgical site was prepared by shaving and cleaning with betadine and alcohol. A 1.5-2 cm incision was made along the mid-line of the scalp, followed by retraction of the scalp and overlying fascia from the skull. A sterile head post was then implanted using dental cement.

### Intrinsic signal imaging

Intrinsic signal imaging (ISI) was done 3 days after headbar surgery to guide later virus injection around the C2 barrel column, and repeated immediately before water restriction started to further confirm the C2 barrel column location in the cranial window. All whiskers except the C2 whisker were trimmed before ISI. To identify the C2 barrel column, the C2 whisker was stimulated using a Piezo stimulator when mice were under light isoflurane anesthesia (0.8-1.0%). Comparisons of acetylcholine dynamics inside the C2 column versus surround were based on column-sized hand-drawn ROIs centered on the ISI hotspot.

### Cranial window and virus injection surgeries

#### Somatosensory task

In all mice, AAV9-hSyn-GRAB-ACh3.0 (Addgene Plasmid #121922 https://www.addgene.org/121922/) was injected from 1x aliquots during the cranial window installation. A glass capillary (Wiretrol® II, Drummond) was pulled to 15-20μm in tip diameter using a micropipette puller (Model P-97, Sutter instrument), and the tip was beveled to about 30 degrees. The glass window was 2x2 mm glass hand fused to 3x3 glass (both 0.13-0.17 mm thickness) with ultraviolet curing glue (Norland optical adhesive 61, Norland Inc.).

Before each surgery, a rimadyl tablet was given 0.5mg/tablet 24 hours prior and Buprenorphine-SR was injected subcutaneously at 0.5 mg/kg immediately before. A 2x2 mm square of skull whose center was the identified C2 whisker barrel region was removed. Virus was backfilled into a pipette of mineral oil (M5904, Sigma-Aldrich). We injected 400nl virus into the identified C2 barrel column through a single injection site over 4 min at 300μm below pia, withdrawing after an additional two minutes. The exposed brain region was then covered with the homemade glass window. Targeting of the C2 column was confirmed via ISI one week after cranial window surgery, before which water restriction commenced.

#### Auditory task

A cranial window was created over the right auditory cortex. The skin over the skull was removed, and part of the temporalis muscle was dissected bluntly to determine the location for the craniotomy. A 3-mm diameter window was opened using a handheld dental drill (Osada Exl-M-40).

In all mice, AAV9-hSyn-GRAB-ACh3.0 (Vigene Biosciences Catalog # YL 10002-AV9) was injected as 1x aliquots during the cranial window installation, using a 5 Microliter Hamilton Syringe (Model 85 RN, Hamilton Company).

We gradually injected 500 nl of the virus into the auditory cortex at a single injection site for approximately 5 minutes, waited an additional 5 minutes, and then withdrew the syringe. The exposed brain region was then covered with the a cranial window, which consisted of two, stacked 3 mm diameter cover glass that were hand-fused to a 5 mm diameter glass (thickness of 0.13-0.17 mm) using ultraviolet curing glue (Norland optical adhesive 71, Norland Inc.). Dental acrylic cement was applied to secure the glass windows to the surrounding bone.

### Water Restriction

Mice were given 1mL of water per day and provided food *ad libitum*. Mice weights stabilized around or above 80% of initial weight. On training days, mice received 0.2∼0.6 mL of water on the rig and were provided supplement water post training, adding up to 1 mL. If the weight fell below 80% of the initial weight, additional water was provided until they returned above 80% of initial weight.

### Behavioral task and training

#### Somatosensory task

Mice were trained in a whisker-guided Go/No-go localization discrimination task^58^. During training, a smooth black pole with 0.6 mm diameter (a plunger for glass capillary, Wiretrol® II, painted with black lubricant, industrial graphite dry lubricant, the B’laster Corp.) was vertically presented at two positions using a pneumatic slide (Festo), with the posterior position rewarded (Go trials) and anterior position non-rewarded (No-go trials). The pole came into touch range within 100 ms of pole motion onset. To promote active whisking to solve the task, we jittered the pole position by 0-2mm along the anterior-posterior axis. Mice used a single whisker (C2) to discriminate positions. Mice indicated their decision through licking or withholding licking during the answer period according to pole position. Licks in the 1 second sampling period were ignored. On Hit trials, mice received water rewards on the first lick in the 1 second answer period. On False Alarm trials, based on each mouse’s learning process, each lick during the answer period re-triggered a timeout that lasted 0-4 seconds. Miss and Correct Rejection trials were unpunished. The behavioral task was controlled by MATLAB-based BControl software (C. Brody, Princeton University).

We trained mice in a stepwise manner. First, we associated the timing of cue/pole with reward to let mice learn that water can come out of the lick port and the water is temporally associated with a pole presentation. Mice usually learned this association in a few minutes. Then we introduced Go trials only to let the mice learn the trial structure, which usually took 1-2 sessions for the mice to achieve high performance in go trials. After the mice were able to get over 10 Hit trials in a row, we introduced No-go trials. During early training, we adjusted the No-go trials probability and time-out time on False Alarm trials to help mice learn, settling on 50% No-go probability and 0s timeout once mice did not get discouraged and stop licking after a series of misses. The threshold for expert performance was defined as a >70% correct rate (Hit trials + Correct Rejection trials/ All trials) continuously for 3 sessions. After the animals became experts, we trimmed the animal’s last whisker (C2 whisker) to test whether the mice learned the task in a whisker-dependent manner.

#### Auditory task

The auditory task (Fig. 5E-H) was the same as previously described ^59^. In short, mice were head-fixed on a wheel and learned to lick for sugar water reward to report detection of an auditory signal stimulus. Each trial consisted of three consecutive intervals: (i) the ‘noise’ or tone cloud interval, (ii) the ‘signal’ (temporal coherence) interval, and (iii) the inter-trial-interval (ITI). The duration of the noise interval was randomly drawn before each trial from an exponential distribution with mean of 5 seconds; this was done to ensure a flat hazard function for signal start time. Randomly drawn noise durations greater than 11 s were set to 11 s. The duration of the ITI was uniformly distributed between 2 and 3 s. Correct-go responses (hits) were followed by either 2 or 12 mL of sugar water (depending on block number; see de Gee et al., bioRxiv, 2022 for the behavioral and pupillometric correlates of changes in reward expectation). Incorrect-go responses (false alarms) terminated the trial and were followed by a 14 s timeout with the same ITI-stimulus.

### Whisker motion acquisition and analysis

Backlit whisker motion video was acquired with a CMOS camera (Basler acA800-500um), StreamPix Software (NorPix Inc.) at 311Hz through telecentric lens (0.09X½” GoldTL™ #58-259, Edmund optics), a CMOS camera (Basler acA800-500um). to record whisker motion. Camera frames were triggered and synced by BControl and Ardunio. We tracked whisker position with the Janelia Whisker Tracker (https://www.janelia.org/open-science/whisk-whisker-tracking ^60^). The fur was masked to improve tracking quality. The whisker’s azimuthal angle was calculated at the intersection of the mask and whisker trace. Whisking midpoint, amplitude and phase was decomposed from this angle using the Hilbert transform, as described in Cheung et al., 2019 ^54^.

### Two-photon microscopy

#### Somatosensory task

The two-photon microscope (Neurolabware) used a galvanometer scanner (6215H, Cambridge Technology), Pockels cell (350-105-02 KD∗P EO modulator, Conoptics), a resonant scanner (CRS8, Cambridge Technology), an objective (W Plan-Apochromat 20×/1.0, Zeiss), a GaAsP photomultiplier tube (H10770B-40, Hamamatsu), and a 510 nm emission filter (FF01-510/84-50, Semrock). We used an 80 MHz tunable laser at 940 nm (Insight DS+, Spectra-Physics) for GRAB-Ach3.0 excitation. Imaging was continuous throughout each session. The scope was controlled by a Matlab-based software Scanbox with custom modifications. Imaging frequency was 15.44 frames/s on the size of the FOV (512 × 796). Spatial resolution was 1.4 μm per pixel.

#### Auditory task

In vivo GRAB-ACh3.0 excitation imaging was conducted in the right auditory cortex using a fast resonant scanning system (ThorLabs rotating Bergamo). The imaging frame rate was set at 15 Hz. Excitation was achieved with a Ti–sapphire laser (Insight DS+, Spectra Physics) tuned to 930 nm, with a ×16 (0.8 NA, Nikon) objective. Imaging was continuous throughout each session. Scan Image (Vidrio) was used to control the imaging system. The imaging data were synchronized with sound stimulation, pupil videos, licking, and wheel motion using custom Labview software.

### Data analysis and statistics

All imaging data were processed in Matlab. We excluded the first 100s for each session due to non-linear photodynamics which stabilized after 100 seconds of continuous excitation scanning (SFig. 2).

Baseline for ΔF/F calculation: In Figure 2B we used the mean fluorescence intensity of the sample session as baseline. To analyze trial based acetylcholine fluorescence changes, in Figure 2C-2E, Figure 3D, and Figure 6F we used the first 16 frames (∼1s) after trial start as baseline. To calculate acetylcholine release triggered by trial cues, in Figure 3A and 3B we used the last 16 frames (-1 to 0s) before stimuli (pole onset and pole withdrawal) as baseline). The acetylcholine fluorescence started to increase around 500 ms before first lick (Figure 4D). To prevent this signal from leaking into the baseline measurement, in Figure 4A, 4D, 5A, 5E, SFigure 4B, and 6A we used frames 32-17 (-2 to -1s) before first lick as baseline. In Figure 4E and 6H, the acetylcholine response to the first lick was calculated by using the mean fluorescent intensity of the one second window immediately following the first lick. In Figure 4F, the acetylcholine transient duration was defined as the time duration between the first lick and the bottom of the first trough of the acetylcholine response. Statistical comparisons were made using paired T-Test corrected for multiple comparisons.

GLM: Over the full dataset for each animal (7 animals, 3 early and 3 expert sessions for each animal), we trained a separate model for each session. We built a design matrix and response matrix for fitting the model, aiming to predict the fluorescence change (dff) value at time t from a set of input parameters leading up to that time t. The design matrix is a matrix of each of these input parameters for each time t that we want to predict. The 9 input parameters are First lick times (the first lick that happens in a trial), Other lick times (all the licks in the trial except the first lick), Reward (water dispense time), Pole in time (pole initiates moving in time), Pole out time (pole initiates moving out time), Amplitude, Midpoint, Phase (3 whisking variables are defined as described in Cheung et al., 2019 ^54^), and Fluorescence history. And we chose the window size and time lags to construct the design matrix. These can be set independently for each input. In the analysis for Figure 7, we used a time lag of 0 for all inputs except Fluorescence history, which is 10 frames. We used a window size of 45 frames for all inputs except for Fluorescence history, which is 5 frames. That is to say that for each time point t we are using the input data history from 45 frames (5 frames for Fluorescence history) before t all the way up to t itself. We did important preprocessing of the data prior to building the design matrix. For continuous data (e.g. whisking amplitude) we normalize the traces between 0 and 1. For data represented as time points (e.g. lick times) we vectorized and binarized the data -i.e. create a vector of zeros with length of all frames in that session. Then we matched the lick times to a frmae time and changed the 0 to 1 at that index. With that we then built a General Linear Model (GLM) for each session. This involves taking a subset of the dataset (first 1/8th of the session) to train the GLM on, fitting the GLM model on the training data, using the fitted GLM to predict the dff value for the full dataset for that session. GLM training used Matlab version of glmnet.

## Supplemental figure legends

**Supplemental Figure 1.**
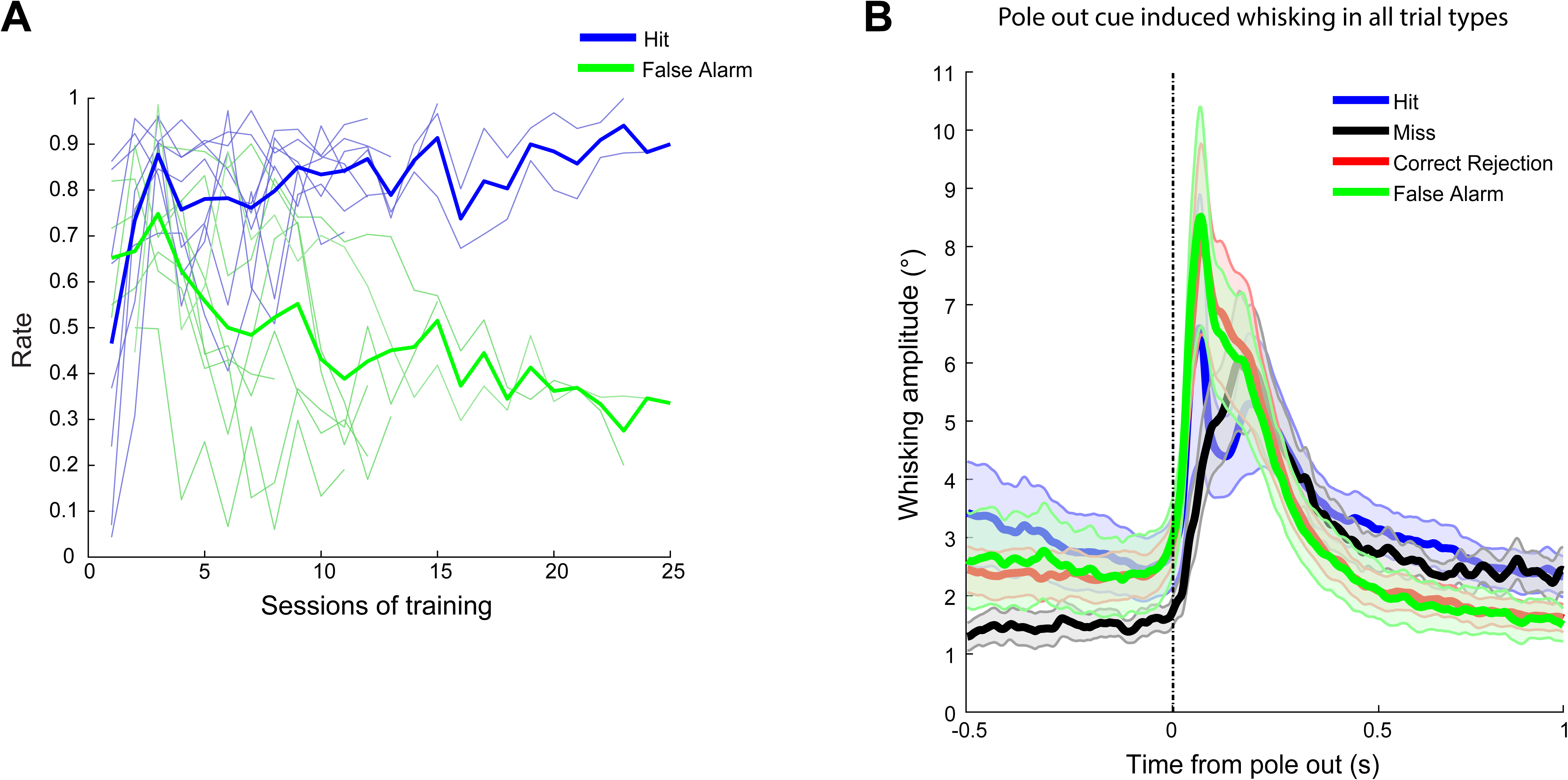
Pole out cue induced whisking in all trial types. **A.** Average Hit rate (blue) and False Alarm rate (green) of 8 mice across training sessions. Individual mouse Hit rate in light blue, False Alarm rate in light green. **B.** Average whisking amplitude aligned to pole out cue by trial outcome, 3 expert sessions / mouse.

**Supplemental Figure 2.**
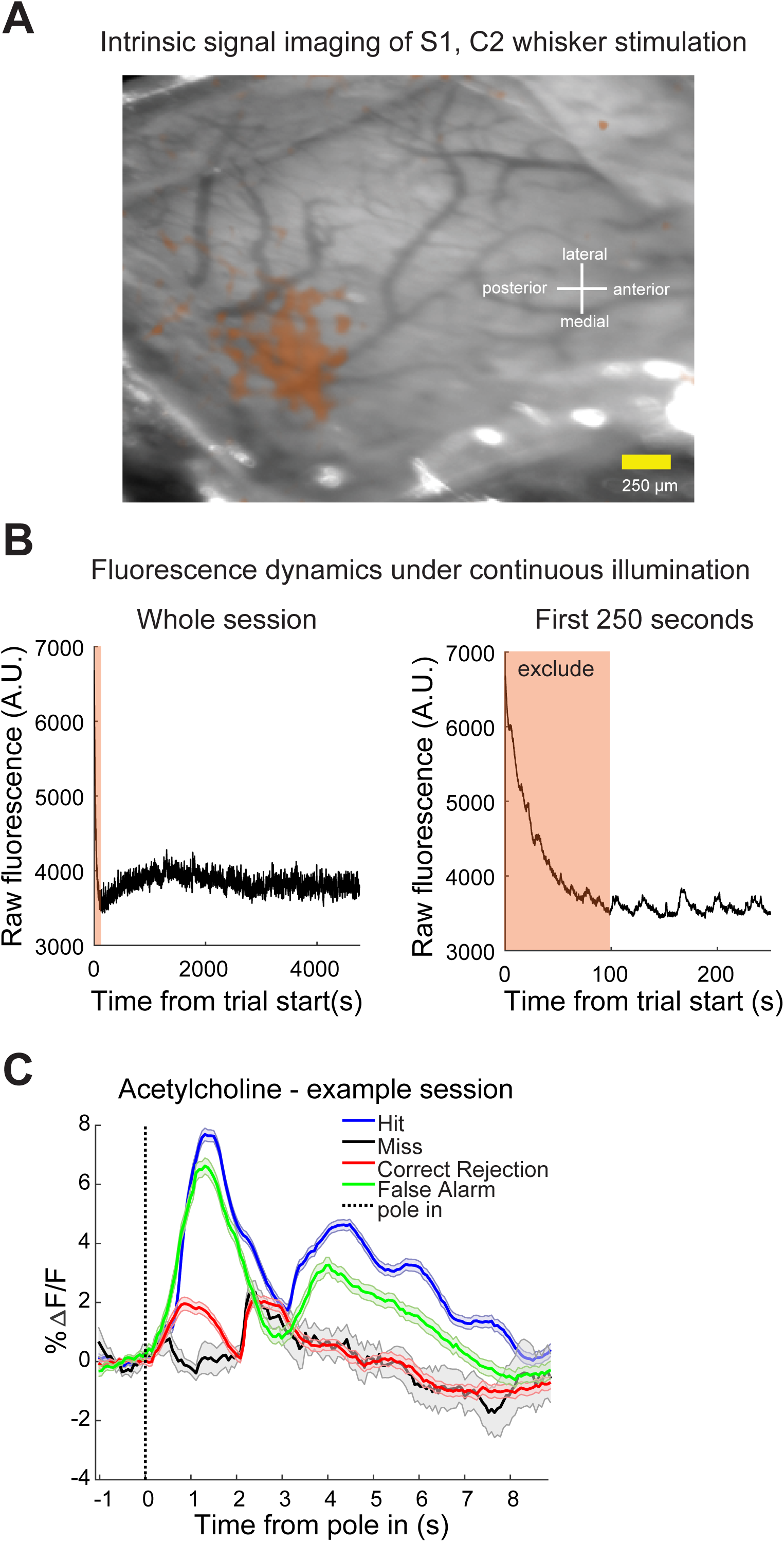
Technical details of targeting and imaging. **A.**Intrinsic signal imaging result showing C2 barrel column location through the cranial window. **B.**Left: Raw fluorescence trace of an example session. The colored area is excluded from analysis due to non-stationary photodynamcis. Right: Zoom of same. **C.** Acetylcholine fluorescence changes averaged by trial types, same session as Figure 2C.

**Supplemental Figure 3.**
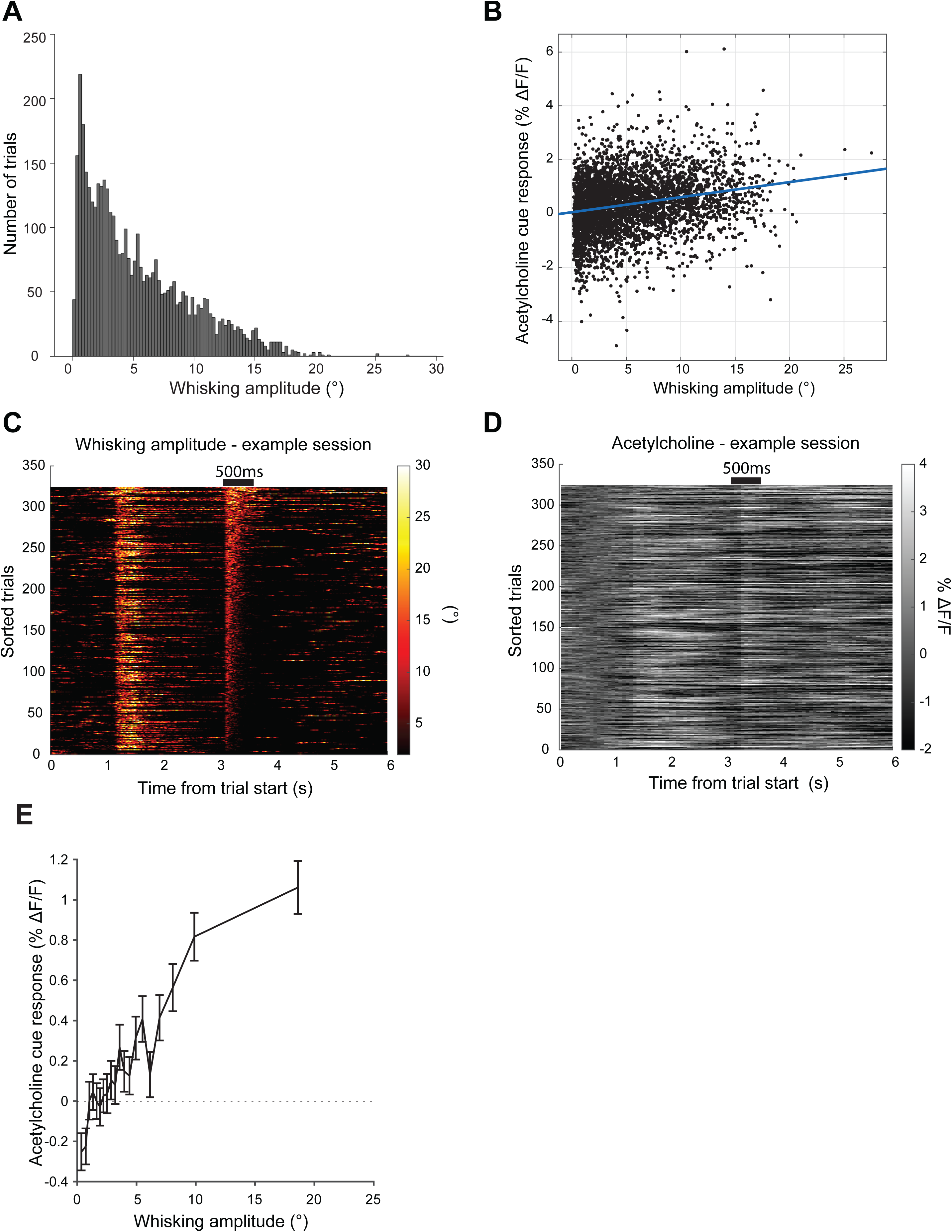
Whisking drives acetylcholine release in S1. **A.** Mean whisking amplitude of 1s post pole in cue. **B.** Mean acetylcholine change vs. mean whisking amplitude over 1s following pole in cue. **C.** Whisking amplitude heatmap sorted by the mean whisking amplitude of 0-500ms post pole withdrawal cue. No lick trials pooled from 3 expert sessions, one mouse. **D.** Acetylcholine release signal heatmap sorted by the same criteria as C. **E.** Mean acetylcholine release signal of 500ms post pole withdrawal cue vs. mean whisking amplitude of 500ms post pole withdrawal cue. No lick trials pooled from 3 early and 3 expert sessions, 8 mice.

**Supplemental Figure 4.**
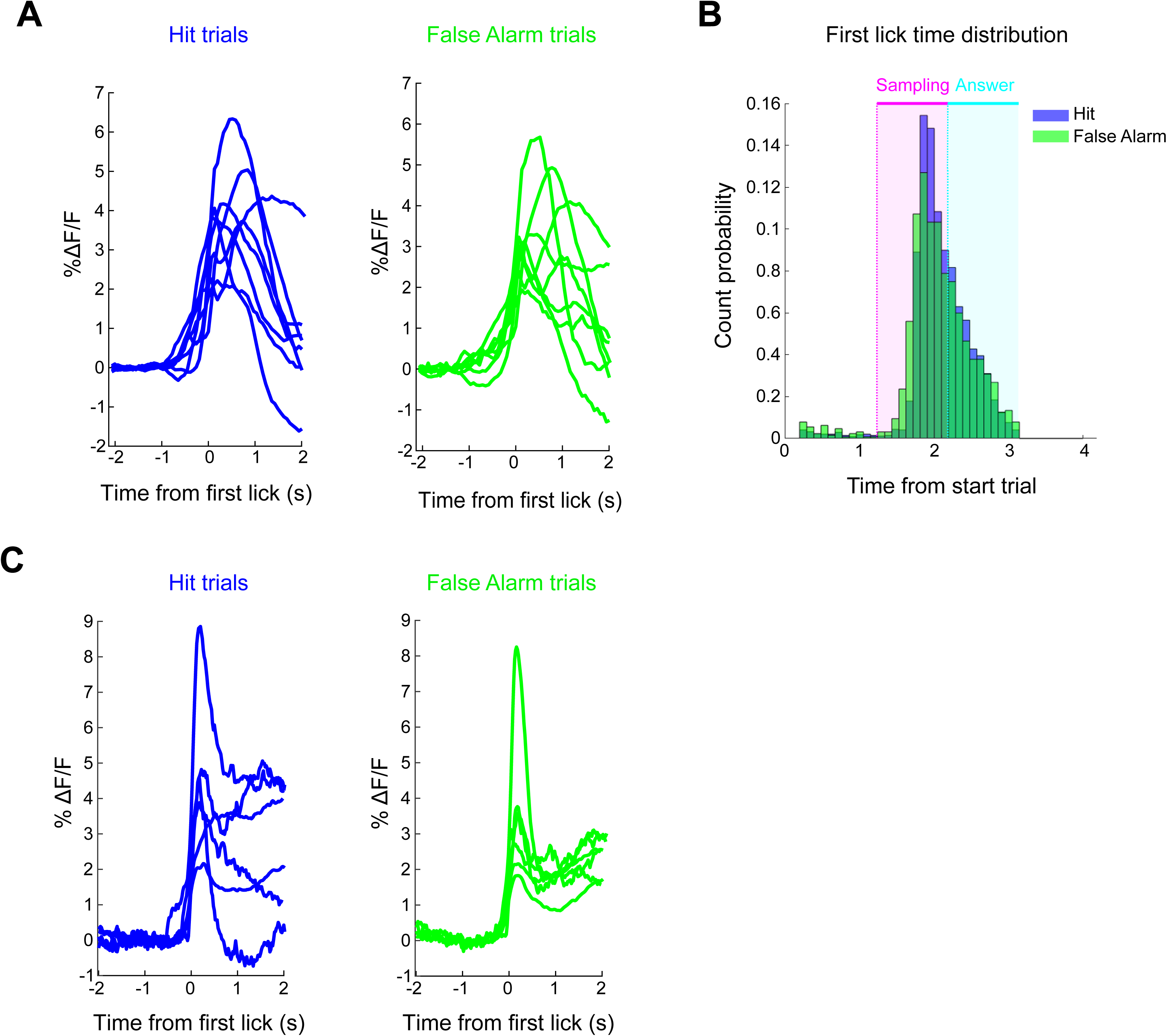
First lick time and first lick related acetylcholine release comparison between Hit and False Alarm trials. **A.** Mean acetylcholine fluorescence change aligned to first lick from Hit trials (left) and False Alarm trials (right) for each mouse (n=8 mice), somatosensory task. **B.** First lick time distribution for Hit (blue) and False Alarm (green) trials. 3 expert sessions, 8 mice. **C.** Mean acetylcholine fluorescence change aligned to first lick from Hit trials (left) and False Alarm trials (right) for each mouse (n=6 mice), Auditory task.

**Supplemental Figure 5.**
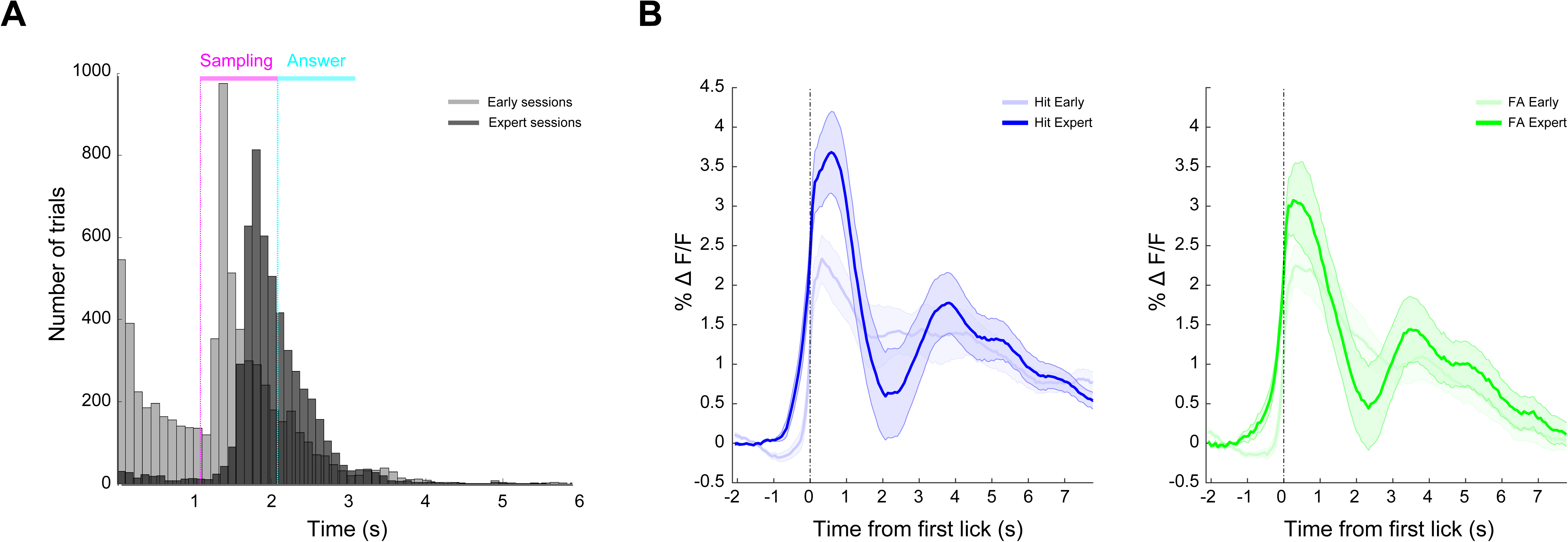
First lick time and first lick related acetylcholine release comparison between early and expert sessions. **A.** First lick time distribution for early (gray) and expert (black) sessions. 3 sessions per condition, 8 mice. **B.** Left, Hit trials average acetylcholine fluorescence change for early (light blue) and expert (blue) sessions. Right, same for False Alarm trials. 3 sessions per condition, 8 mice.

## References

1. Parikh, Vinay, et al. "Prefrontal acetylcholine release controls cue detection on multiple timescales." Neuron 56.1 (2007): 141–154.

2. Zhang, Hao, Shih-Chieh Lin, and Miguel AL Nicolelis. "A distinctive subpopulation of medial septal slow-firing neurons promote hippocampal activation and theta oscillations." Journal of neurophysiology 106.5 (2011): 2749–2763.

3. Kim, Jae-Hyun, et al. "Selectivity of neuromodulatory projections from the basal forebrain and locus ceruleus to primary sensory cortices." Journal of Neuroscience 36.19 (2016): 5314–5327.

4. Reimer, Jacob, et al. "Pupil fluctuations track rapid changes in adrenergic and cholinergic activity in cortex." Nature communications 7.1 (2016): 1–7.

5. Donoghue, John P., and Kristen L. Carroll. "Cholinergic modulation of sensory responses in rat primary somatic sensory cortex." Brain research 408.1–2 (1987): 367–371.

6. Tremblay, Nicole, Richard A. Warren, and Robert W. Dykes. "Electrophysiological studies of acetylcholine and the role of the basal forebrain in the somatosensory cortex of the cat. II. Cortical neurons excited by somatic stimuli." Journal of neurophysiology 64.4 (1990): 1212–1222.

7. Hars, Bernard, et al. "Basal forebrain stimulation facilitates tone-evoked responses in the auditory cortex of awake rat." Neuroscience 56.1 (1993): 61–74.

8. Goard, Michael, and Yang Dan. "Basal forebrain activation enhances cortical coding of natural scenes." Nature neuroscience 12.11 (2009): 1444–1449.

9. Chubykin, Alexander A., et al. "A cholinergic mechanism for reward timing within primary visual cortex." Neuron 77.4 (2013): 723–735.

10. Liu, Cheng-Hang, et al. "Selective activation of a putative reinforcement signal conditions cued interval timing in primary visual cortex." Current Biology 25.12 (2015): 1551–1561.

11. Crouse, Richard B., et al. "Acetylcholine is released in the basolateral amygdala in response to predictors of reward and enhances the learning of cue-reward contingency." Elife 9 (2020): e57335.

12. Sturgill, James Fitzhugh, et al. "Basal forebrain-derived acetylcholine encodes valence-free reinforcement prediction error." BioRxiv (2020): 2020–02.

13. Kilgard, Michael P., and Michael M. Merzenich. "Cortical map reorganization enabled by nucleus basalis activity." Science 279.5357 (1998): 1714–1718.

14. Froemke, Robert C., et al. "Long-term modification of cortical synapses improves sensory perception." Nature neuroscience 16.1 (2013): 79–88.

15. Jiang, Li, et al. "Cholinergic signaling controls conditioned fear behaviors and enhances plasticity of cortical-amygdala circuits." Neuron 90.5 (2016): 1057–1070.

16. Guo, Wei, Blaise Robert, and Daniel B. Polley. "The cholinergic basal forebrain links auditory stimuli with delayed reinforcement to support learning." Neuron 103.6 (2019): 1164–1177.

17. Hasselmo, Michael E., and Martin Sarter. "Modes and models of forebrain cholinergic neuromodulation of cognition." Neuropsychopharmacology 36.1 (2011): 52–73.

18. Lee, Seung-Hee, and Yang Dan. "Neuromodulation of brain states." Neuron 76.1 (2012): 209–222.

19. Zaborszky, Laszlo, et al. "Neurons in the basal forebrain project to the cortex in a complex topographic organization that reflects corticocortical connectivity patterns: an experimental study based on retrograde tracing and 3D reconstruction." Cerebral cortex 25.1 (2015): 118–137.

20. Li, Xiangning, et al. "Generation of a whole-brain atlas for the cholinergic system and mesoscopic projectome analysis of basal forebrain cholinergic neurons." Proceedings of the National Academy of Sciences 115.2 (2018): 415–420.

21. Robert, Blaise, et al. "A functional topography within the cholinergic basal forebrain for processing sensory cues associated with reward and punishment." bioRxiv (2021).

22. Inglis, F. M., and H. C. Fibiger. "Increases in hippocampal and frontal cortical acetylcholine release associated with presentation of sensory stimuli." Neuroscience 66.1 (1995): 81–86.

23. Eggermann, Emmanuel, et al. "Cholinergic signals in mouse barrel cortex during active whisker sensing." Cell reports 9.5 (2014): 1654–1660.

24. Chaves-Coira, Irene, et al. "Basal forebrain nuclei display distinct projecting pathways and functional circuits to sensory primary and prefrontal cortices in the rat." Frontiers in neuroanatomy 12 (2018): 69.

25. Herrero, Jose L., et al. "Acetylcholine contributes through muscarinic receptors to attentional modulation in V1." Nature 454.7208 (2008): 1110–1114.

26. Pinto, Lucas, et al. "Fast modulation of visual perception by basal forebrain cholinergic neurons." Nature neuroscience 16.12 (2013): 1857–1863.

27. Hangya, Balázs, et al. "Central cholinergic neurons are rapidly recruited by reinforcement feedback." Cell 162.5 (2015): 1155–1168.

28. Harrison, Thomas C., et al. "Calcium imaging of basal forebrain activity during innate and learned behaviors." Frontiers in neural circuits 10 (2016): 36.

29. Richardson, Russell T., and M. R. DeLong. "Context-dependent responses of primate nucleus basalis neurons in a go/no-go task." Journal of Neuroscience 10.8 (1990): 2528–2540.

30. Clarke, Samuel, and Jay A. Trowill. "Sniffing and motivated behavior in the rat." Physiology & behavior 6.1 (1971): 49–52.

31. Bindra, D., and J. F. Campbell. "Motivational effects of rewarding intracranial stimulation." Nature 215.5099 (1967): 375–376.

32. Jing, Miao, et al. "An optimized acetylcholine sensor for monitoring in vivo cholinergic activity." Nature methods 17.11 (2020): 1139–1146.

33. Lohani, Sweyta, et al. "Dual color mesoscopic imaging reveals spatiotemporally heterogeneous coordination of cholinergic and neocortical activity." bioRxiv (2020).

34. Kimchi, Eyal Y., et al. "Reward contingency gates selective cholinergic suppression of amygdala neurons." bioRxiv (2022).

35. Bollu, Tejapratap, et al. "Cortex-dependent corrections as the tongue reaches for and misses targets." Nature 594.7861 (2021): 82–87.

36. Kim, Jinho, et al. "Behavioral and Neural Bases of Tactile Shape Discrimination Learning in Head-Fixed Mice." Neuron 108.5 (2020): 953–967.

37. Guo, Zengcai V., et al. "Procedures for behavioral experiments in head-fixed mice." PloS one 9.2 (2014): e88678.

38. Moore, Jeffrey D., David Kleinfeld, and Fan Wang. "How the brainstem controls orofacial behaviors comprised of rhythmic actions." Trends in neurosciences 37.7 (2014): 370–380.

39. Klinkenberg, Inge, Anke Sambeth, and Arjan Blokland. "Acetylcholine and attention." Behavioural brain research 221.2 (2011): 430–442.

40. McGaughy, Jill, et al. "Selective behavioral and neurochemical effects of cholinergic lesions produced by intrabasalis infusions of 192 IgG-saporin on attentional performance in a five-choice serial reaction time task." Journal of Neuroscience 22.5 (2002): 1905–1913.

41. Kuchibhotla, Kishore V., et al. "Parallel processing by cortical inhibition enables context-dependent behavior." Nature neuroscience 20.1 (2017): 62–71.

42. O’Connor, Daniel H., et al. "Neural activity in barrel cortex underlying vibrissa-based object localization in mice." Neuron 67.6 (2010): 1048–1061.

43. Gasselin, Celia, et al. "Cell-type-specific nicotinic input disinhibits mouse barrel cortex during active sensing." Neuron 109.5 (2021): 778–787.

44. Ramamurthy, Deepa L., et al. "VIP interneurons in sensory cortex encode sensory and action signals but not direct reward signals." Current Biology 33.16 (2023): 3398–3408.

45. Shadlen, Michael N., and William T. Newsome. "Motion perception: seeing and deciding." Proceedings of the national academy of sciences 93.2 (1996): 628–633.

46. Letzkus, Johannes J., et al. "A disinhibitory microcircuit for associative fear learning in the auditory cortex." Nature 480.7377 (2011): 331–335.

47. Urban-Ciecko, Joanna, et al. "Precisely timed nicotinic activation drives SST inhibition in neocortical circuits." Neuron 97.3 (2018): 611–625.

48. Lee, Soohyun, et al. "A disinhibitory circuit mediates motor integration in the somatosensory cortex." Nature neuroscience 16.11 (2013): 1662–1670.

49. Gasselin, Célia, et al. "Cell-type-specific nicotinic input disinhibits mouse barrel cortex during active sensing." Neuron 109.5 (2021): 778–787.

50. Takahashi, Naoya, et al. "Active cortical dendrites modulate perception." Science 354.6319 (2016): 1587–1590.

51. Takahashi, Naoya, et al. "Active dendritic currents gate descending cortical outputs in perception." Nature Neuroscience 23.10 (2020): 1277–1285.

52. Lacefield, Clay O., et al. "Reinforcement learning recruits somata and apical dendrites across layers of primary sensory cortex." Cell reports 26.8 (2019): 2000–2008.

53. Cheung, Jonathan Andrew, et al. "Independent representations of self-motion and object location in barrel cortex output." PLoS Biology 18.11 (2020): e3000882.

54. Cheung, Jonathan, et al. "The sensorimotor basis of whisker-guided anteroposterior object localization in head-fixed mice." Current Biology 29.18 (2019): 3029–3040.

55. Williams, Leena E., and Anthony Holtmaat. "Higher-order thalamocortical inputs gate synaptic long-term potentiation via disinhibition." Neuron 101.1 (2019): 91–102.

56. Yao, Zizhen, et al. "A taxonomy of transcriptomic cell types across the isocortex and hippocampal formation." Cell 184.12 (2021): 3222–3241.

57. Bloem, Bernard, et al. "Topographic mapping between basal forebrain cholinergic neurons and the medial prefrontal cortex in mice." Journal of Neuroscience 34.49 (2014): 16234–16246.

58. O’Connor, Daniel H., et al. "Vibrissa-based object localization in head-fixed mice." Journal of Neuroscience 30.5 (2010): 1947–1967.

59. de Gee, J. W., et al. "Mice regulate their attentional intensity and arousal to exploit increases in task utility." bioRxiv (2022).

60. Clack, Nathan G., et al. "Automated tracking of whiskers in videos of head fixed rodents." PLoS computational biology 8.7 (2012): e1002591.

